# Tumor microenvironment impaired T cell antigen recognition and function were restored by Lovastatin therapy

**DOI:** 10.1101/2022.09.11.507496

**Authors:** Zhou Yuan, Meghan J. O’Melia, Kaitao Li, Jintian Lyu, Aaron M. Rosado, Valencia E. Watson, Amir Hossein Kazemipour Ashkezari, Fangyuan Zhou, Vikash Kansal, Brendan Kinney, Stefano Travaglino, Larissa O. Doudy, Richard K. Noel, Samuel N. Lucas, Steven Lingyang Kong, Prithiviraj Jothikumar, Nathan A. Rohner, Margaret P. Manspeaker, David M. Francis, Ke Bai, Chenghao Ge, Muaz N. Rushdi, Loice Chingozha, Samuel Ruipérez-Campillo, Ning Jiang, Hang Lu, Nicole C. Schmitt, Susan N. Thomas, Cheng Zhu

## Abstract

CD8^+^ T cells underpin effective anti-tumor immune responses in melanoma; however, their functions are attenuated due to various immunosuppressive factors in the tumor microenvironment (TME), resulting in disease progression. T cell function is elicited by the T cell receptor (TCR), which recognizes antigen peptide-major histocompatibility complex (pMHC) expressed on tumor cells via direct physical contact, i.e., two-dimensional (2D) interaction. TCR–pMHC 2D affinity plays a central role in antigen recognition and discrimination, and is sensitive to both the conditions of the T cell and the microenvironment in which it resides. Herein, we demonstrate that CD8^+^ T cells residing in TME have lower 2D TCR–pMHC bimolecular affinity and TCR–pMHC–CD8 trimolecular avidity, pull fewer TCR–pMHC bonds by endogenous forces, flux lower level of intracellular calcium in response to antigen stimulation, exhibit impaired *in vivo* activation, and show diminished anti-tumor effector function. These detrimental effects are localized in the tumor and tumor draining lymph node (TdLN), and affect both antigen-inexperienced and antigen-experienced CD8^+^ T cells irrespective of their TCR specificities. These findings implicate impaired antigen recognition as a mechanism of T cell dysfunction in the TME.

## Introduction

Spontaneous development of CD8^+^ T cell immunity against melanomas is associated with improved clinical outcomes (*1, 2*). Immunotherapy therefore has a high potential for the treatment of advanced melanoma, which is among the common cancer types with poor clinical outcomes at advanced disease stages (*3*). The protective effects of effector CD8^+^ T cells are often inhibited within the tumor microenvironment (TME), however, resulting in unchecked tumor growth (*4–8*). To this end, therapeutic immune checkpoint blockade (ICB) of programmed cell death 1 (PD-1) signaling, thought to work both by expanding the pool of pre-existing stem-like CD8^+^ T cells (*9*) and reinvigorating exhausted tumor-infiltrating lymphocytes (TILs) from immune-suppression (*10, 11*), has emerged as a successful treatment that induces durable objective responses in 25-30% of patients with advanced melanoma (*12, 13*). ICB therefore represents a highly promising therapy class to address the high and increasing mortality associated with melanoma. However, the success of ICB therapies relies on the existence of robust, pre-existing CD8^+^ T cell immunity (*14, 15*). Elucidating the mechanisms suppressing both the development and function of CD8^+^ T cell anti-tumor immunity against endogenously generated tumor neoantigens during disease progression or through therapeutic vaccination can therefore help develop strategies to improve the success of melanoma immunotherapy.

T cell anti-tumor immunity is initiated by antigen recognition of the T cell receptor (TCR) via interaction with peptide-major histocompatibility complex (pMHC). TCR–pMHC affinity is an important driver of CD8^+^ T cell anti-tumor immunity, which is conventionally measured by the surface plasmon resonance (SPR) technology using aglycosylated soluble proteins of TCR ectodomain produced by *E. coli* or pMHC-tetramer staining of T cells (*16*). However, *in vivo* TCR–pMHC interaction requires direct physical contact between the T cell and the tumor cell as the two molecules reside on the respective cell surfaces and interact across the junctional gap between the two opposing cell membranes.

We (*17–22*) and others (*23*) have shown that TCR–pMHC binding affinity measured *in situ* at the T cell surface better correlates with T cell function. Such a measurement is termed two-dimensional (2D) affinity because it has a unit of area, in contrast to the 3D affinity measured by SPR that has a unit of volume. Similar to 3D affinity whose value may depend on pH or chemical composition of the medium, the value of 2D affinity can be regulated by pharmacological perturbations of the cellular environment *in vitro* (*17, 24*). Importantly, *ex vivo* measurements found differential 2D affinities of monoclonal TCR on membranes of T cells isolated from splenic red pulp vs white pulp, which are well correlated with differential transcriptomic changes, differential effector functions, and differential developmental fates of these cells *in vivo* (*25*). Remarkably, it was found that 2D affinity correlated with these phenotypes even in the cases where the surface markers for cell subsets did not (*25*), highlighting the predictive power of our 2D affinity measurement.

Using the B16F10 melanoma model most widely used for the preclinical investigation of immunotherapeutic resistance, we reveal that the 2D affinity for and spontaneous pulling of TCR on its cognate pMHC by CD8^+^ T cells is impaired by the TME in which the T cells reside, and that the lowered 2D TCR–pMHC affinity predicts suppressed effector functions. These findings suggest altered TCR antigen recognition as a mechanism of immune evasion in melanoma, independent of the inhibitory effects of immune checkpoint receptors, whose consideration may help develop diagnostic and immunotherapeutic approaches to manage advanced disease.

## Results

### TME suppresses antigen-induced expansion and function of CD8^+^ T cells *in vivo*

The influence of the TME on CD8^+^ T cell immunity was explored using a murine tumor model (Fig. 1A) and a synthetic antigen system in which lymph-draining nanoparticles (NPs, Fig. 1B) were disulfide-linked to the H2-K^b^ presented immunodominant peptide of ovalbumin (OVA) with a C-terminal cysteine, CSIINFEKL, (CSIINFEKL-NP or Ag-NP, Fig. 1C) that result in dose-dependent cross-presentation in *in vitro* splenocyte cultures (Fig. 1D) (*26*). In this model, C57BL/6 mice were implanted in the lateral dorsal skin with a primary B16F10 melanoma at day (d) −8 (Fig. 1A, left). At d-1 (7 days post primary tumor implantation), Ag-NPs were administered intratumorally (i.t.) into the primary B16F10 melanoma (Tumor-bearing) (Fig. 1A, left). Alternatively, Ag-NPs were administered into the skin of tumor-naïve animals (Naïve) (Fig. 1A, right) at d-1. In both schema (Fig. 1A), mice were implanted with OVA-expressing B16F10 (B16F10-OVA) melanoma cells at d0 (one day post Ag-NP administration) in either the dorsal skin contralateral to the B16F10 melanoma or naïve skin contralateral to the Ag-NP injection site in mice of the Tumor-bearing and Naïve cohorts, respectively. At d2, the Ag-NP injection site (the primary B16F10 melanoma for the Tumor-bearing animals or skin for the Naïve animals) was resected. In so doing, we could compare the influences on the growth of the OVA-expressing secondary tumor and animal survival by the microenvironment in which the anti-OVA immunity had been elicited by Ag-NP administration into naïve skin vs the primary tumor. As Ag-NPs result in priming of CD8^+^ T cells both at the local injection site as well as its draining lymph nodes (dLNs) (*26*) due to these NPs unique lymph-draining behaviors (*27–32*), this system allowed us to compare microenvironmental effects on CD8^+^ T cells exerted both locally at the site of injection and within lymph nodes (LNs) draining the naïve skin vs tumor after synthetic antigen administration without animals succumbing to the primary tumor. Control of the B16F10-OVA secondary melanoma was found to be less effective when Ag-NP treatment was administered into the primary B16F10 melanoma of the tumor-bearing animals compared to the skin of tumor-naïve animals, reflected in both larger tumor volumes 14 days post injection (Fig. 1E) and shortened survival (Fig. 1F). These data indicate that endogenous CD8^+^ T cells primed by Ag-NP within the TME elicit less effective anti-tumor responses than those primed by Ag-NP administration in the tumor-free (naïve) skin.

**Fig. 1.**
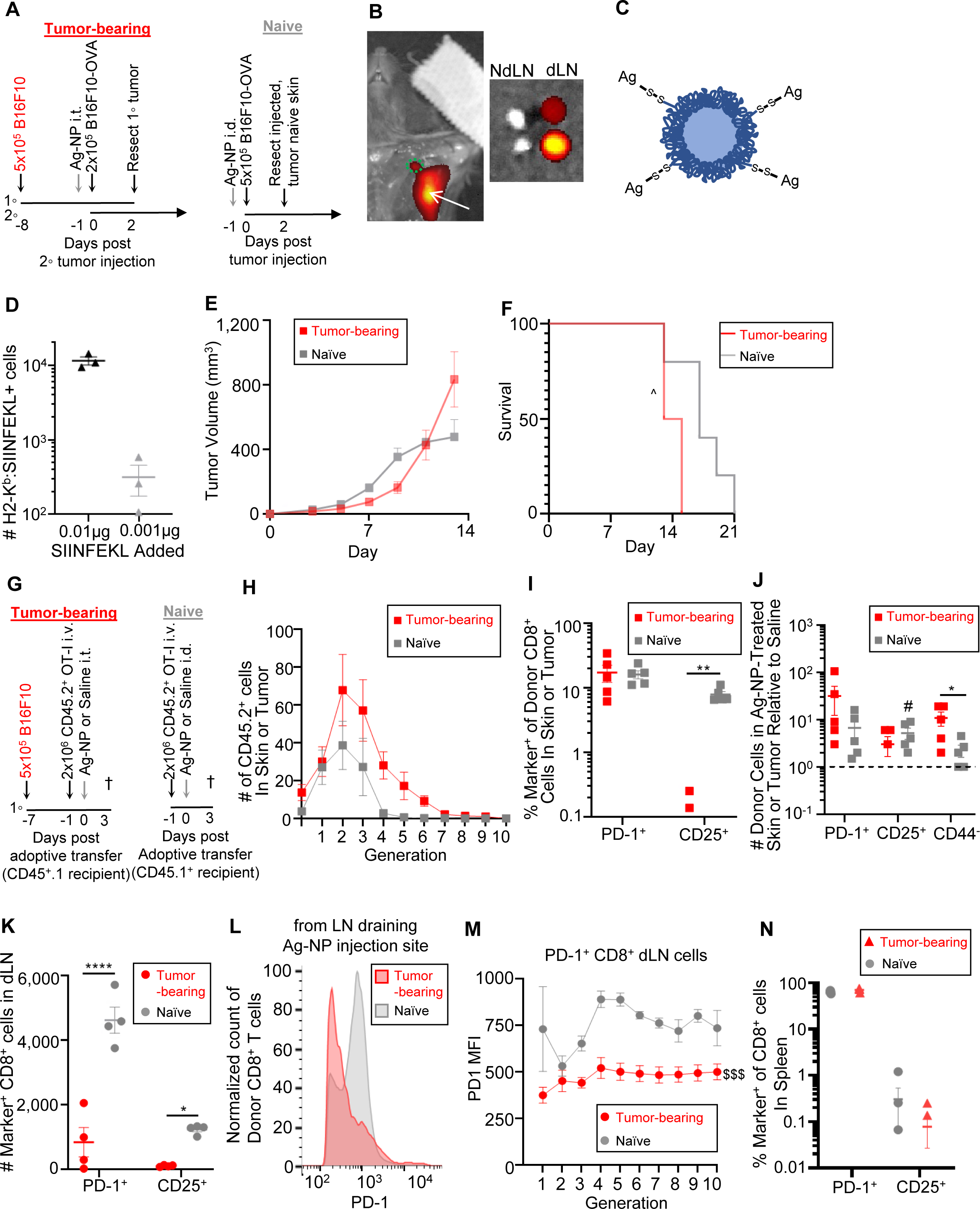
Functionally impaired antigen-specific CD8^+^ T cell responses in the TME. **(A)** Animal models used in (*B*)-(*F*). *Left*, B16F10 melanoma cells were intradermally injected into the lateral dorsal skin of C57BL/6 mice at d-8. Ag-NPs were administered at d-1 into the tumor. *Right*, Ag-NPs were administered into the tumor-naïve lateral dorsal skin at d-1. *Both*, at d0, B16F10-OVA melanoma cells were injected into the contralateral dorsal skin. Primary melanomas were resected at d2 and tumor growth as well as animal survival were monitored. **(B)** IVIS image of AF647 signal 24 h post AF647-labelled NP administration i.d. into the lateral dorsal skin. *Left*, NP injection site (white arrow) and dLN (green circle). *Right*, dLN and NdLN after excision. **(C)** Schematic of the Ag-NPs. **(D)** Number of CD3^-^ cells positively stained by a mAb against SIINFEKL:H2-K^b^ (clone 25D1.16) 24 h after *in vitro* coincubation of SIINFEKL with splenocytes. Each point represents a technical replicate. Mean ± SD (lines). **(E, F)** Mean ± SEM (n=5) of size of contralateral (secondary) tumor (*E*) and animal survival (*F*) after Ag-NP challenge in the tumor (red) or naïve skin (gray), corresponding to the model cohorts depicted in **(G)** Animal model used in (*H*)-(*N*). *Left*, B16F10 melanoma cells were injected into CD45.1^+^ C57BL/6 recipient mice at d-7. *Left and right*, CD8^+^ T cells isolated from the spleens of CD45.2^+^ OT-I donor mice were adoptive transferred at d-1 and Ag-NPs or saline was administered i.t. into B16F10 bearing animals (Tumor-bearing) or the naïve skin (Naïve) at d0. At d3 the donor cells were harvested from recipient mice and analyzed. **(H)** Number of CD45.2^+^ OT-I cells in each proliferative generation in response to *in vivo* challenge with Ag-NP in the tumor (red) of the tumor-bearing animals and the naïve skin of the control animals (gray). Data represents mean ± SEM. n=5. **(I-K)** Comparison of frequency (*I*) and number (*J*, *K*) of indicated phenotypic marker expressing CD45.2^+^ populations between cells harvested from the tumor (*i, j*) or the LNs draining the tumor (*K*) of the tumor-bearing animals (red) and from naïve skin (*I, J*) or the LNs draining the naïve skin (*K*) of the control animals (gray) after *in vivo* Ag challenge. Each point represents individual animal. Data represent mean ± SEM. n=5. **(L, M)** Comparison of anti-PD-1 mAb fluorescence intensity (FI) profile using representative histogram from donor cells from an individual animal from each group (*L*) and mean ± SEM (n=5) of mean FI (MFI) of PD-1^+^subpopulation in each proliferative generation (*M*) between CD45.2^+^ cells harvested from the LNs draining the tumor of the tumor-bearing animals (red) or the naïve skin of the control animals (gray). **(N)** Frequency of PD-1^+^ and CD25^+^ cells over CD45.2^+^ population in the spleen of tumor-bearing (red) or tumor-lacking (gray) animals after *in vivo* Ag challenge. Each point represents individual animal. Data represent mean ± SEM. n=5. “*” indicates significance by two-way ANOVA with Tukey’s post-hoc test; “#” indicates significance compared to theoretical value of 1.0 by one-sample t-test; “^” indicates significance by Log-rank (Mantel-Cox) test; $ indicates significance by repeat measures ANOVA; Tumor growth and survival experiments in (*E*)-(*F*) are representative of two independent experiments.

To further test the above contention, we compared the effects of the melanoma TME on the antigen-specific activation and expansion of carboxyfluorescien succinimidyl ester (CFSE) labelled CD8^+^ OT-I T cells transferred to C57BL/6 animals in response to the OVA antigen delivered by Ag-NP (Fig. 1G). Tumor-bearing CD45.1^+^ recipient mice were implanted with B16F10 melanomas at d-7, received adoptive transfer of CD8^+^ T cells isolated from CD45.2^+^ OT-I donor mice at d-1, were injected i.t. with either Ag-NPs or saline on d0, and were sacrificed at d3 to recover the donor CD8^+^ T cells to measure their quantity and quality (Fig. 1G, left). For comparison, donor CD8^+^ T cells were likewise assessed after recovered from tumor-naïve CD45.1^+^recipient mice (comprising the Naïve cohort) that received adoptive transfer of CD8^+^ T cells isolated from CD45.2^+^ OT-I donor mice d-1, were intradermally (i.d.) administered Ag-NPs or saline one day later (d0), and were sacrificed at d3 (three days post injection of Ag-NP) (Fig. 1G, right). No statistically significant differences in the overall extent of proliferation by donor CD8^+^ T cells localized within the tissue site of injection were seen between the Tumor-bearing and - Naïve groups challenged with Ag-NP (Fig. 1H), in line with reports that antigenic priming induces a programmed proliferation response in CD8^+^ T cells (*33*). The frequency and relative number of donor CD8^+^ T cells expressing PD-1, which was increased upon recent antigen experience or chronic antigen stimulation (*34*), was similar in Tumor-bearing and -Naïve recipient animals in response to Ag-NP challenge (Fig. 1I-J). However, the frequency of CD8^+^ donor cells expressing the activation marker CD25 in response to Ag-NP treatment was significantly lower in tumors of Tumor-bearing animals than the naïve skin injection site of Naïve animals (Fig. 1I). To account for differences in lymphocyte homing to tumors vs skin (*35*), the overall numbers of donor CD8^+^ T cells in tumors of Tumor-bearing animals and skin of Naïve animals were respectively normalized to those measured in animals from each group that had been challenged with saline instead of Ag-NPs. With this method that accounts for intrinsic microenvironmental differences in total levels of lymphocyte accumulation in the tissue site of analysis, CD25-expressing donor CD8^+^ T cell numbers that resulted specifically from Ag-NP challenge were found to increase in the skin of saline injected Naïve animals but not in tumors of Tumor-bearing animals. Ag-NP challenge likewise increased the number of CD44^-^ donor cells in tumors of Tumor-bearing animals relative to the skin of Naïve animals (Fig. 1J). These data indicate that the TME locally suppresses overall levels of antigen-specific CD8^+^ T cell activation in response to antigen challenge.

Given the known role of LNs draining the tumor (TdLNs) in mediating adaptive immunity and response to therapy (*26, 36, 37*), responses to Ag challenge of donor CD8^+^ T cells recovered from TdLNs of Tumor-bearing animals were compared to those from LNs draining the Ag-NP-injected skin of Naïve animals. The total numbers of PD-1- and CD25-expressing donor CD8^+^ T cells were significantly lower in the TdLNs of Tumor-bearing animals than those draining the Ag-NP injected skin of Naïve animals (Fig. 1K), in contrast to the lack of local changes within the tumor of Tumor-bearing animals relative to the skin injection site of Naïve animals. Moreover, PD-1 expression was significantly lower in donor CD8^+^ T cells recovered from TdLNs of Tumor-bearing animals intratumorally challenged with Ag-NPs than LNs draining the Ag-NP injected skin of Naïve animals (Fig. 1L), an effect observed in both early and late proliferative generations (Fig. 1M). However, statistically indistinguishable levels of PD-1 or CD25 were observed on donor CD8^+^ T cells recovered from spleens of Tumor-bearing and Naïve recipient animals after Ag challenge (Fig. 1N). Collectively, these results demonstrate that CD8^+^ T cells within the TME and TdLNs, but not in spleen, exhibit less robust response to antigenic priming *in vivo*.

### Activation by and TCR binding to antigen pMHC is suppressed in CD8^+^ T cells recovered from TME and TdLNs

The diminished responses of CD8^+^ T cells to antigen priming within tumor and TdLNs (collectively referred to as TME) were further explored using CD45.1^+^ C57BL/6 mice that were tumor naïve (Naïve) or had been implanted B16F10 melanoma (Tumor) at d-7, received adoptive transfer of CD45.2^+^ CD8^+^ OT-I T cells at d0, and sacrificed at d3 to harvest donor cells from the tumor, skin or dLNs (Fig. 2A). Donor CD8^+^ T cells were stimulated *in vitro* with SIINFEKL, and responses were measured by intracellular cytokine staining, finding lower frequency of activation marker CD69-expressing cells harvested from the tumor than from the skin of naïve animals (Fig. 2B). Furthermore, a higher frequency of donor cells from the tumor than skin expressed tumor necrosis factor (TNF) α, whereas similar frequencies of donor cells from the same two sites expressed interferon (IFN) γ, interleukin (IL) 2, and granzyme (GzmB) (Fig. 2B). Moreover, significantly lower frequency of donor CD8^+^ T cells from TdLNs of Tumor-bearing animals than skin-dLNs of Naïve animals expressed CD69 and PD-1, whereas similar frequencies of donor cells from the same two sites expressed TNF-α, IFN-γ, IL-2, and GzmB (Fig. 2C). By comparison, statistically indistinguishable numbers of donor CD8^+^ T cells were found from the spleens of tumor-bearing and -naive recipient mice that expressed CD69, PD-1, TNF-α, IFN-γ, IL-2, and GzmB (Fig. 2D). These results corroborate data generated in response to *in vivo* antigen encounter and show that microenvironmental factors derived from the tumor, which are present in TdLNs but not spleens in this model (*37, 38*), suppresses CD8^+^ T cell activation induced by antigen recognition.

**Fig. 2.**
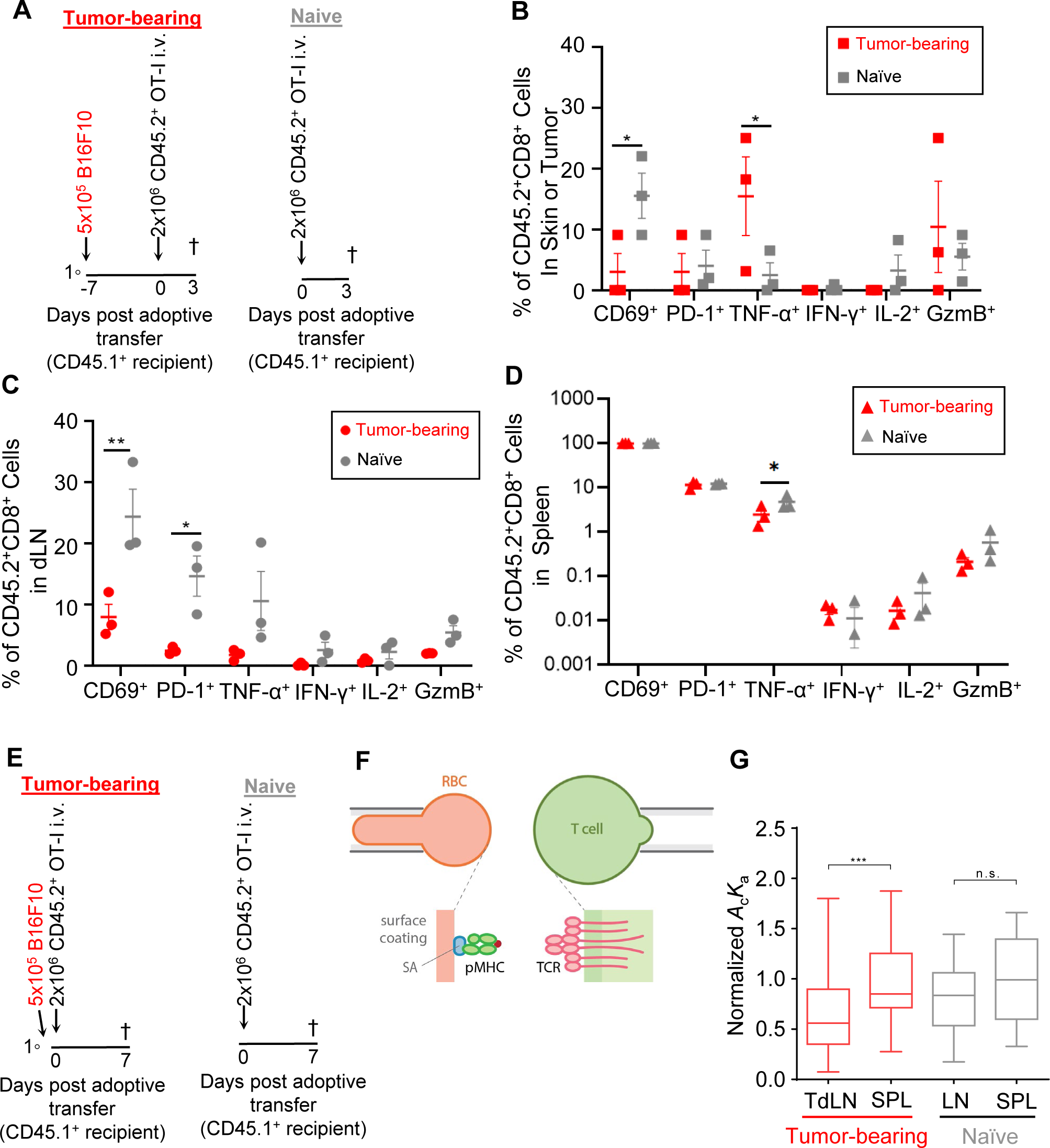
Lowered T cell activation response and TCR–pMHC 2D affinity of CD8^+^ T cells recovered from tumors and TdLNs. **(A)** Animal models used in (*B*)-(*D*). *Left*, CD45.1^+^ C57BL/6J recipients were implanted with B16F10 tumor on d-7. *Left and right*, splenic CD8^+^ T cells from CD45.2^+^ OT-I mice adoptively transferred into recipient animals d0 and donor cells were harvested d3. **(B-D)** Activation and function in response to *ex vivo* SIINFEKL challenge of CD45.2^+^ donor cells harvested from tumor vs naïve skin (*B*), tumor-vs naïve-dLNs (*C*), and spleens (*D*) of tumor-bearing (red) vs -naïve (gray) animals were analyzed by for the indicated phenotypical markers. Each point represents data from one individual animal. Data represents mean ± SEM. Representative results of two independent experiments with n≥3. **(E)** Animal model used in (*G*). CD45.1^+^ C57BL/6J recipients that were tumor-naïve (*right*) or had just been implanted with B16F10 tumors (*left*) were adoptively transferred with splenic CD8^+^ T cells from CD45.2^+^ OT-I mice on d0. At d7, CD45.2^+^ donor cells were isolated from recipients’ TdLNs and spleen for binding analysis. **(F)** Schematic of micropipette adhesion frequency (MAF) assay. A micropipette-aspirated RBC coated with SINNFEKL:H2-K^b^α3A2 (*left*) was driven to repeatedly contact a donor CD8^+^ T cell expressing OT-I TCR, aspirated by an opposing micropipette (*right*). Adhesion mediated by TCR–pMHC interaction, if occurred, was detected visually by RBC elongation. An adhesion frequency (*P*_a_ = # of adhesions/# of total contacts) was evaluated from 50 times of repeated 2-s contacts between a RBC–T cell pair from which an effective 2D affinity *A*_c_*K*_a_ = - ln(1 – *P*_a_)/(*m*TCR*m*pMHC) was calculated. Here *A*_c_ is the constant contact area and *m*TCR and *m*pMHC are the respective densities of TCR and pMHC measured separately by FACS. **(G)** Comparison of TCR–pMHC 2D affinities measured from indicated conditions. 2D affinity of each compartment was divided by the mean *A*_c_*K*_a_ value of T cells from the spleen of the same animal (to normalize inter-animal variations). Results shown in (*B*)-(*D*) were from n= 3-5 mice; “*” indicates significance by two-way ANOVA with Tukey’s post-hoc test. Data in (*G*) are presented as mean (line) ± 75/25% (boxes) and max/min (whiskers) of pooled measurements from two independent sets of experiments in order to test sufficient number (24–31) of T cells per condition to ensure reliable statistical comparisons (Mann-Whitney test). ***p<0.001, ns indicates p≥0.05.

Antigen recognition is initiated by TCR–pMHC interaction that occurs at the cross-junctional space spanning the gap between the membranes of the T cell and the tumor cell, i.e., 2D binding, which could be regulated by cellular and cytokine environment (*25*). The effects of the TdLN on TCR–pMHC 2D affinity on donor CD45.2^+^ CD8^+^ OT-I T cells recovered 7 days post adoptive transfer into CD45.1^+^ recipient animals co-implanted with (Tumor, Fig. 2E, left) or without (Naïve, Fig. 2e, right) B16F10 tumors were evaluated by the micropipette adhesion frequency (MAF) assay (*17, 39*). MAF employs a human red blood cell (RBC) coated with pMHC as a surrogate tumor cell to interact with the T cell, which were respectively aspirated by two opposing micropipettes (Fig. 2F). The ultrasoft RBC acts as a force transducer to detect TCR– pMHC binding from RBC elongation after its retraction from contact with the T cell. Instead of H2-K^b^, a mutant H2-K^b^α3A2 was used to present the OVA peptide SIINFEKL, which replaced the mouse α3 domain of H2-K^b^ by that of human HLA-A2, in order to abolish any mouse CD8– MHC interaction (*17*). Each T cell was repeatedly contacted by the RBC 50 times to determine an adhesion frequency from which a 2D affinity was evaluated (*17, 39*). Similar TCR–pMHC 2D affinities were found from donor CD8^+^ OT-I T cells isolated from LNs and spleens of naïve recipient animals; however, significantly lower 2D affinities were found from OT-I CD8^+^ T cells isolated from TdLNs than spleens of the same tumor-bearing recipient animals (Fig. 2G). This shows that the TdLN diminished TCR binding affinity for antigen pMHC, explaining, at least in part, the observed TME suppression of CD8^+^ T cell activation in vivo (Fig. 1E, F, H-M) and in vitro (Fig. 2B, C).

### Characterization of TME suppression of TCR–pMHC and TCR–pMHC–CD8 interactions

To further characterize the localized suppressive effects of tumor and TdLN on TCR antigen recognition, TCR 2D affinity for antigen pMHC was systematically evaluated comparing among CD8^+^ T cells recovered from tumors, TdLNs, non-tumor associated LNs (NdLNs), the spleen and blood. For these experiments, B16F10 cells not expressing the OVA antigen were directly implanted into OT-I mice and the endogenous CD8^+^ T cells were isolated from compartments of interest at various days post tumor implantation for MAF analysis (Fig. 3A). This model was used to overcome the overall low levels of donor cell recovery in the adoptive transfer models (Fig. 1G, 2E), yielding as many as 100-fold CD8^+^ T cells expressing OT-I TCR specific for SIINFEKL:H2-K^b^ from not only lymphoid tissues but also the tumor itself. B16F10 melanoma growth in the two tumor-recipient mouse strains was comparable (Fig. S1A). Lymphoid tissues including TdLNs (Fig. S1B) and spleen also exhibited no gross differences in the frequencies of major immune cell types except for decreased macrophage and increased B lymphocyte frequencies in OT-I animals relative to C57BL/6 animals (Fig. S1C). Immune phenotyping the melanoma-implanted OT-I and C57BL/6 mice over the course of tumor development also revealed no significant differences in the numbers of CD8^+^ T cells infiltrating the tumor, within TdLN or spleen between the OT-I and C57BL/6 mice, except for a mild expansion of CD8^+^ T cells in the spleen of OT-I animals from d7 to d11 post tumor implantation (Fig. S1D). No gross differences were found in antigen activation phenotypes as assessed by the numbers of PD-1 (Fig. S1E) or CD25 (Fig. S1F) expressing CD8^+^ T cells, except for an increase in splenic PD-1^+^CD8^+^ population in OT-I animals at d11. However, the number of CD4^+^ T cells were ∼10-fold lower in OT-I than C57BL/6 animals (Fig. S1G), including CD25^+^Foxp3^+^CD4^+^ regulatory T cells (Tregs) (Fig. S1H), as expected (*40*). The number of myeloid-derived suppressor cells (MDSCs) and PD-1 ligand-1 (PD-L1)-expressing lymphocytes (PD-L1^+^CD45^+^) were equivalent between analyzed tissues of OT-I and C57BL/6 animals throughout tumor progression, except for subtle increases in in spleens or tumors of the OT-I animals on d7-d11 or d11 respectively (Fig. S1I-J). Thus, the abundance and phenotype of CD8^+^ T cells as well as immune suppressive cell subsets are roughly similar for OT-I and C57BL/6 tumor-bearing mice, indicating similar TMEs established in both C57BL/6 and OT-I hosts.

**Fig. 3.**
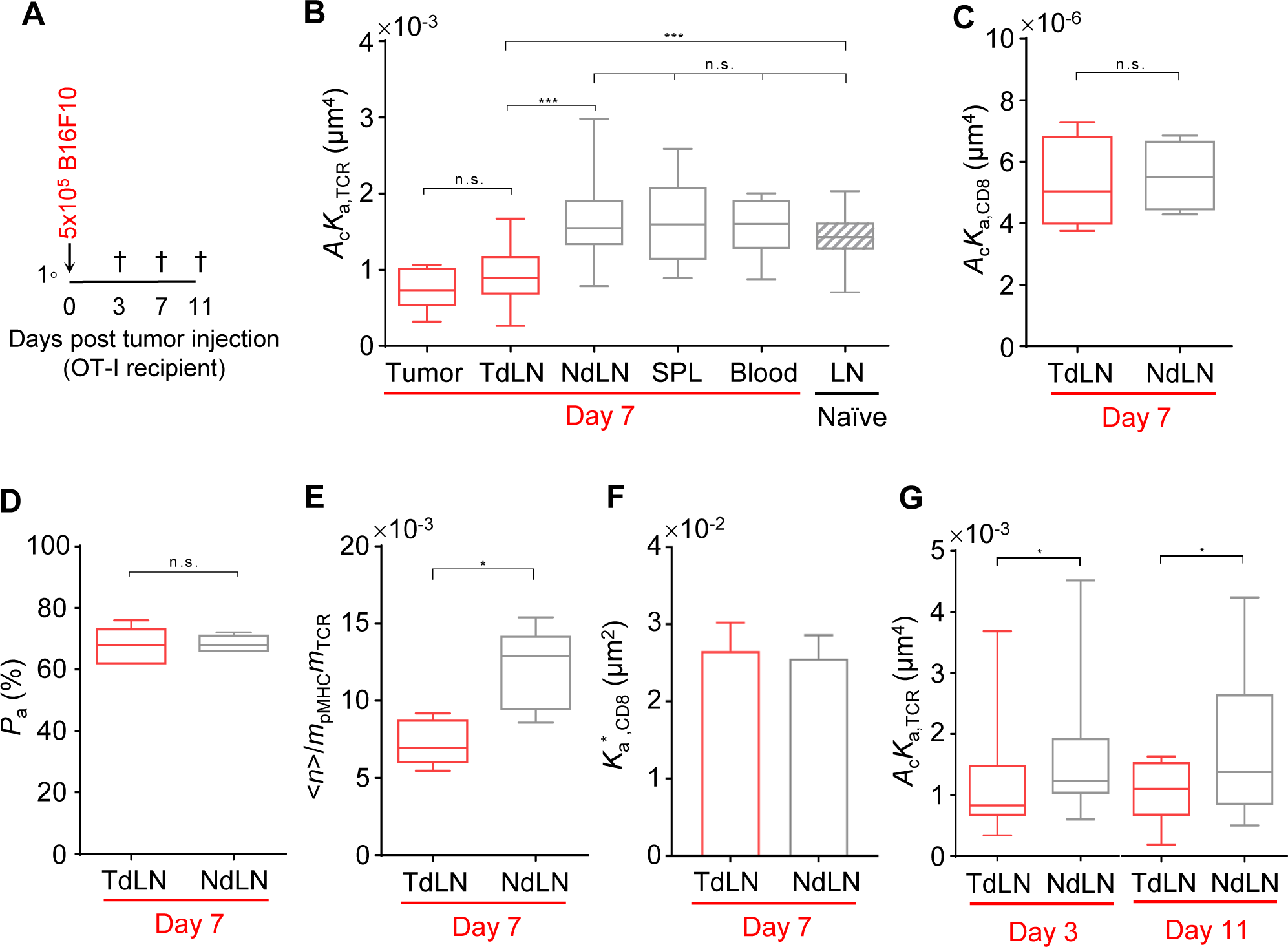
Suppressed TCR antigen recognition by tumor and TdLN-localized CD8^+^ T cells. **(A)** Animal model used in (*B*)-(*G*) in which CD8^+^ T cells were isolated from indicated tissues of B16F10 tumor-bearing OT-I mice at indicated timepoints for analysis. **(B)** Comparison of OT-I TCR–SIINFEKL:H2-K^b^α3A2 2D affinities evaluated using CD8^+^ T cells from the indicated tissues. **(C)** Comparison of CD8–VSV:H2-K^b^ 2D affinities evaluated using CD8^+^ T cells from TdLN and NdLN of the same OT-I mouse. **(D)** Comparison between adhesion frequencies of RBCs bearing anti-TCR antibody and CD8^+^ T cells from TdLN and NdLN of the same OT-I animal. **(E)** Comparison of average numbers of bonds per contact from OT-I CD8^+^ T cells– SIINFEKL:H2-K^b^ interaction, calculated from adhesion frequencies by <*n*> = - ln(1 – *P*_a_), normalized by the densities of the TCR (*m*TCR) and pMHC (*m*pMHC), evaluated using CD8^+^ T cells from TdLN and NdLN. Since <*n*>/(*m*TCR*m*pMHC) is the effective 2D affinity *A*_c_*K*_a,TCR_ of the TCR–pMHC bonds in the absence of CD8 binding, the 1 log higher <*n*>/(*m*TCR*m*pMHC) values observed here (which include not only TCR–pMHC bonds but also TCR–pMHC–CD8 and CD8–MHC bonds) than the corresponding values in (*B*) (which include only TCR–pMHC bonds) indicate that the trimolecular bonds greatly outnumber the two bimolecular bonds. **(F)** Comparison of 2D affinities of CD8 for pre-formed TCR–pMHC complex, *K*_a_*_,CD8_ = [<*n*>/(*A*_c_*K*_a,TCR_*m*TCR*m*pMHC) – 1]/*m*CD8, evaluated using CD8^+^ T cells from TdLN and NdLN of the same OT-1 mouse. The calculation of *K* * used the CD8 site density (*m*CD8) measured from flow cytometry, and the data from (*B*) and (*E*) by subtracting the contribution of the TCR– pMHC bonds from the total bonds. The contribution of CD8–MHC bond was neglected since the CD8–MHC affinity is ∼500-fold smaller than the TCR–pMHC affinity. **(G)** Same as (*B*) except that the experiments were done 3- and 11-days post tumor injection for only two tissue sources. Data are presented as mean (line) ± 75/25% (boxes) and max/min (whiskers) of pooled measurements from 2-6 independent experiments in order to test sufficient number (4–27) of T cells per condition to ensure reliable statistical comparisons (Mann-Whitney test).

The TCR–SIINFEKL:H2-K^b^α3A2 2D affinities were measured by the MAF assay (Fig. 2F) using CD8^+^ T cells isolated from various tissue compartments of OT-I mice. Cells from tumor and TdLNs had much lower affinities than those from NdLN, spleen, and blood (Fig. 3B), agreeing with previous results (Fig. 2G) obtained using the model in which OT-I donor cells were adoptively transferred into melanoma-bearing C57BL/6 recipients and harvested from the TdLN 7 days post tumor implantation (Fig. 2E). Furthermore, CD8^+^ T cells from NdLN, spleen, and blood of tumor-bearing OT-I mice exhibited comparable TCR–pMHC 2D affinities to that of CD8^+^ T cells isolated from LNs of tumor-free OT-I mice (Fig. 3B), suggesting the local nature of this suppressive effect that is limited to the tumor and TdLN. However, MHC–CD8 2D affinities measured using a null pMHC (VSV:H2-K^b^) not recognized by the OT-I TCR showed no compartmental differences (Fig. 3C), isolating the TME-induced defect to the TCR molecule. Moreover, OT-I T cells from TdLN and NdLN generated similar levels of binding frequencies to an anti-TCR Vα2 antibody (Fig. 3D), indicating the antigen-specific nature of this suppressive effect that impairs the ability for the TCR to recognize antigen. Still, when OT-I T cells were tested against SIINFEKL:H2-K^b^ to allow CD8 to interact with the MHC to form TCR–pMHC–CD8 trimolecular bonds (*18, 41*), a decrease in the average number of bonds (

<*n*>) normalized by the densities of TCR (*m*TCR) and pMHC (*m*pMHC) was again observed using CD8^+^ T cells from NdLN to TdLN (Fig. 3E). However, the compartmental differences between NdLN and TdLN disappeared when the 2D affinities of CD8 to bind MHC pre-bound by TCR were compared (*42*) (Fig. 3F). Thus, the impaired TCR–pMHC– CD8 trimolecular interaction can be attributed to a reduced TCR–pMHC 2D affinity rather than the synergy between TCR and CD8 for pMHC binding. Finally, the reduced TCR–pMHC 2D affinity of CD8^+^ T cells from NdLNs to TdLNs was detectable as early as d3 post tumor implantation, and was sustained in OT-1 mice bearing d11 tumors (Fig. 3G). This reduction in TCR–pMHC 2D affinity ranged from 20-40% in d3, d7, and d11 post tumor implantation (Supp Fig. 2A). Thus, antigen recognition by CD8^+^ T cells was persistently suppressed during tumor progression in both tumor and TdLN as signified by the decreased TCR–pMHC bimolecular affinity and TCR–pMHC–CD8 trimolecular avidity. Note that 2D affinity differences were found despite no difference in TCR expression and staining by tetramers with SIINFEKL:H-2K^b^ and SIINFEKL:H-2K^b^α3A2 on T cells isolated from various tissues of tumor-bearing and -naive OT-I mice (Fig. S2B-C), consistent with previous findings that 2D analysis is superior to tetramer analysis (*18, 21, 25*).

Transforming growth factor (TGF) β in TME is known to play pleiotropic roles, including modulating tumor growth and the functions of CD8^+^ T cells (*43–45*). TGF-β has furthermore been shown to be a determinant of the differential TCR–pMHC 2D affinities of CD8^+^ T cells from splenic red pulp and white pulp of virial infected mice (*25*) and to negatively regulate tumor-specific CD8^+^ T cell cytotoxicity (*46*). The effect of inhibiting TGF-β signaling on TCR–pMHC 2D affinity was examined through daily intraperitoneal injection of TGF-β R1 inhibitor SB431542 in OT-I mice bearing B16F10 melanomas (Fig. 4A). In contrast to vehicle controls, the difference in TCR–pMHC 2D affinities between CD8^+^ T cells from TdLNs and NdLNs of TGF-β R1 inhibitor treated animals was no longer observed (Fig. 4B). This implicates the TME-related soluble factor TGF-β in deficient CD8^+^ T cells antigen recognition.

**Fig. 4.**
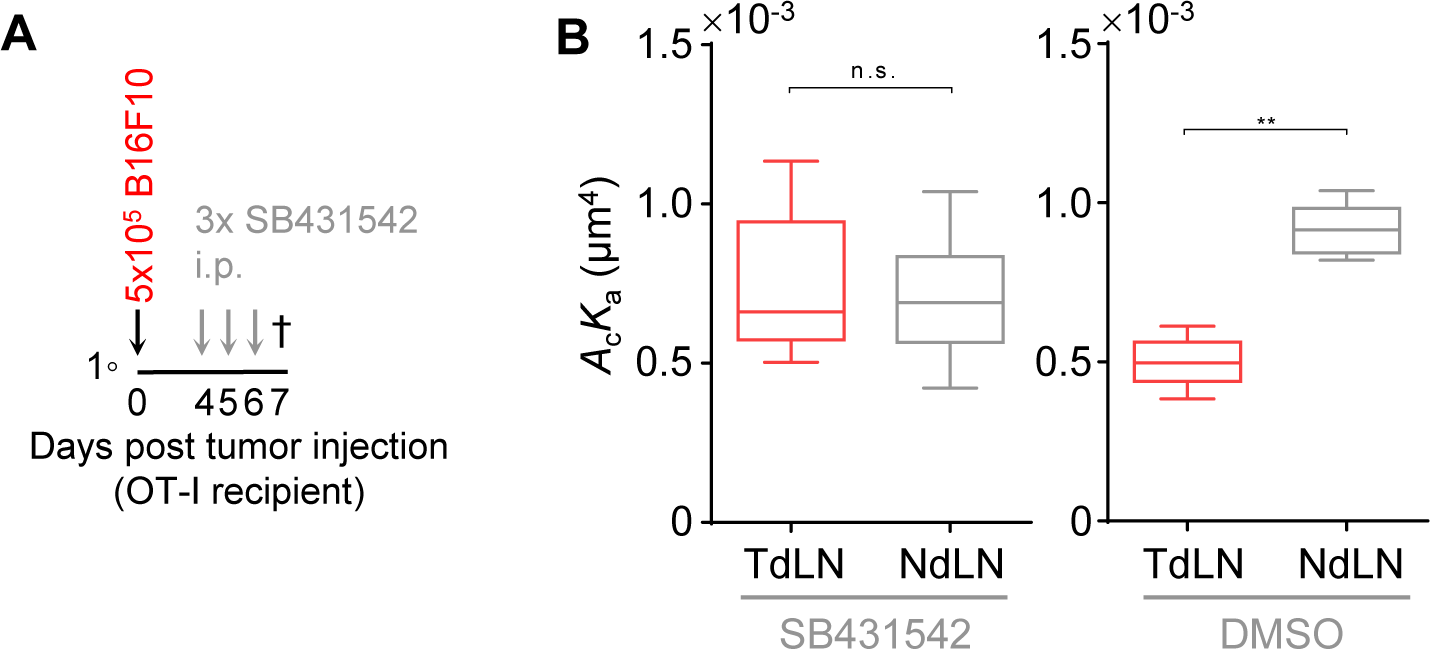
TGF-β inhibition eliminates differential TCR–pMHC 2D affinities between CD8^+^ T cells from TdLNs vs NdLNs. **(A)** Animal model used in (*B*). B16F10 tumor-bearing OT-I mice were treated with TGF-β inhibitor or DMSO as control in d4, d5, and d6 post tumor implantation. **(B)** Comparisons of OT-I TCR–SIINFEKL:H2-K^b^α3A2 2D affinities evaluated using CD8^+^ T cells from TdLN and NdLN of mice treated with TGF-β inhibitor (*left*) or DMSO (*right*). Data are presented and statistically analyzed the same way as Fig. 3.

### TdLN suppresses CD8^+^ T cells pulling forces on antigen pMHC

To elicit anti-tumor immunity, T cells must interact physically with tumor cells. Upon engaging their TCR with (neo)antigenic pMHC, T cells may exert endogenous forces on the TCR–pMHC and TCR–pMHC–CD8 bonds as part of their mechano-sensing and -responsive processes, which has been observed *in vitro* using DNA-based molecular tension probes (MTP) (*47, 48*). MTPs are a class of powerful tools for measuring ligand-induced, signaling-dependent, and actomyosin-powered endogenous forces exerted by the cell on the receptor–ligand bonds (*49–51*). An MTP consists of a fluorophore-quencher pair flanking a DNA hairpin to unfold at a designed tension adjustable by changing the GC contents and length of the DNA strands (*49, 52, 53*) (Fig. 5A). One end presents a ligand or antibody to the cell surface receptor and the other end is linked to the cover glass *via* a gold nanoparticle to further quench the fluorophore (e.g., Cy3B). Forced-unfolding of the DNA-hairpin de-quenches the Cy3B to yield a fluorescent signal, thereby reporting an above-threshold tension exerted on the receptor by the cell (Fig. 5A). The fluorescent intensity is proportional to the number of unfolded DNA hairpins, hence reads out the ability of the TCR and/or CD8 to bind, stay engaged, and be activated by pMHC, and of the T cell to generate and apply above threshold forces on the TCR–pMHC and TCR–pMHC–CD8 bonds, all of which are features of CD8^+^ T cells that we would like to test whether TME would suppress.

**Fig. 5.**
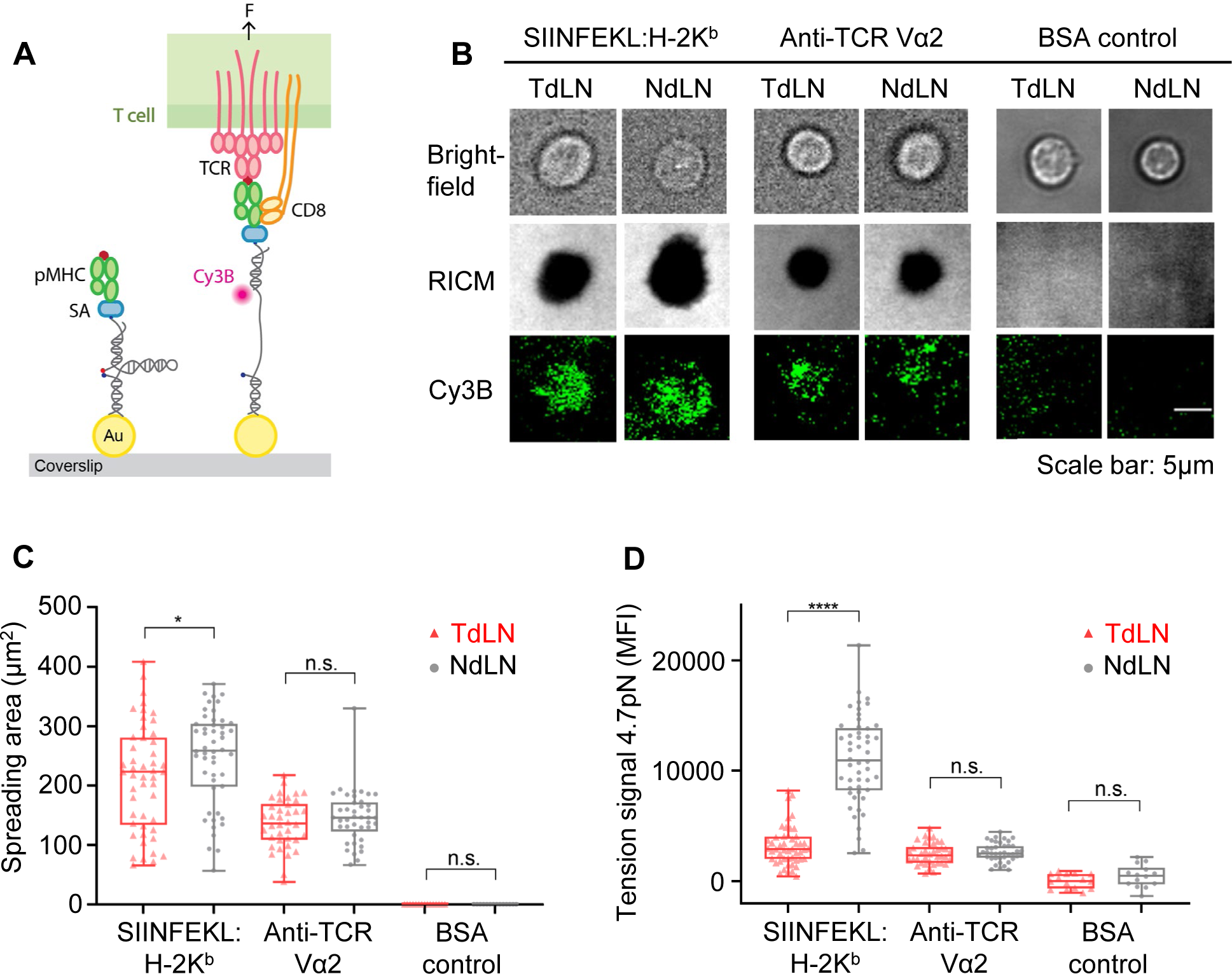
Cell spreading and pulling on antigen pMHC is diminished for CD8^+^ T cells isolated from TdLNs. **(A)** Schematic of DNA-based MTP, consisting of a Cy3B-quencher pair flanking a DNA hairpin to unfold at >4.7 pN forces. One end presents a pMHC or antibody to the TCR and/or CD8 and the other end is linked to the cover glass *via* a gold nanoparticle to further quench the fluorophore. A >4.7 pN force exerted on the pMHC or antibody by the cell unfolds the DNA-hairpin and de-quenches the Cy3B to yield a fluorescent signal. **(B)** Representative images of CD8^+^ T cells from TdLNs or NdLNs of OT-I mice 7 days post tumor implantation placed on glass surfaces functionalized with MTPs tagged by SIINFEKL:H2-K^b^, anti-TCR or BSA viewed 10 min after cell seeding by bright-field (top row), RICM (middle row) and TIRF (bottom row). **(C, D)** Quantification of the spreading area (middle row in *B*) (*C*) and mean fluorescence intensity (MFI) of the force signal (bottom row in *B*) (*D*) using a large number of cells. Points, mean (line) ± 75/25% (boxes) and max/min (whiskers) from a representative experiment that repeated three times. Analyzed CD8^+^ T cells were recovered from TdLNs vs NdLNs of d7 B16F10 melanoma-bearing animals as depicted in Fig. 3a. ****p≤0.0001, *p≤0.05, n.s.: p≥0.05.

CD8^+^ T cells from TdLNs or NdLNs of OT-I mice bearing d7 B16F10 tumors (Fig. 3A) were placed on glass surface functionalized with SIINFEKL:H2-K^b^ tagged by MTP of a 4.7 pN threshold force. Twenty min post cell seeding, the area of T cell spreading was measured by reflection interference contrast microscopy (RICM) and the average fluorescence intensity over that area was measured by total internal reflection fluorescence (TIRF) microscopy (Fig. 5B). As shown by the quantification obtained from a large number of cells, CD8^+^ T cells from TdLNs were found to spread less (Fig. 5C) and pull on less pMHC molecules with >4.7 pN forces (Fig. 5D) than cells from NdLNs. Remarkably, no such differential spreading areas and force signals between TdLN and NdLN CD8^+^ T cells were observed when the MTP tag was changed from the SIINFEKL:H2-K^b^ ligand to an anti-TCR antibody (Fig. 5C, D). The antigen-specific nature of these two differential readouts correlates with that of the differential TCR interactions with pMHC and lack of difference of TCR interactions with the same anti-TCR antibody (Fig. 3B, D), supporting a hypothetical relationship between the TdLN impaired TCR 2D affinity and the TdLN impaired T cell activation, spreading as well as force generation, transmission, and exertion on pMHC.

### TdLN suppresses calcium signaling of CD8^+^ T cells stimulated by antigen pMHC

To directly test the hypothesis that TdLN impairs antigen-induced T cell activation, OVA pMHC-triggered calcium signaling of CD8^+^ T cells from TdLNs and NdLNs of OT-I mice bearing d7 B16F10 melanomas (Fig. 3A) were compared. T cells were loaded with a calcium indicator Fluo-4 and a microfluidic cell trap array device was used to place them on polydimethylsiloxane (PDMS) surfaces functionalized with SIINFEKL:H2-K^b^ or bovine serum albumin (BSA) as control (Fig. 6A, *top*) (*54*). Calcium signals of cells imaged in the field of view were recorded continuously for 10 min in real-time at the single-cell level by their fluorescence intensity (Fig. 6A, *bottom*). For each cell, the moment when a cell settled into the cell trap was defined as “t = 0”. The fluorescence intensity time-courses for different cells were realigned to their respective starting times, which were displayed on a heatmap as columns and sorted in ascending cumulative calcium intensity across rows, clearing showing that T cells generated calcium responses to pMHC but not BSA (Fig. 6B).

**Fig. 6.**
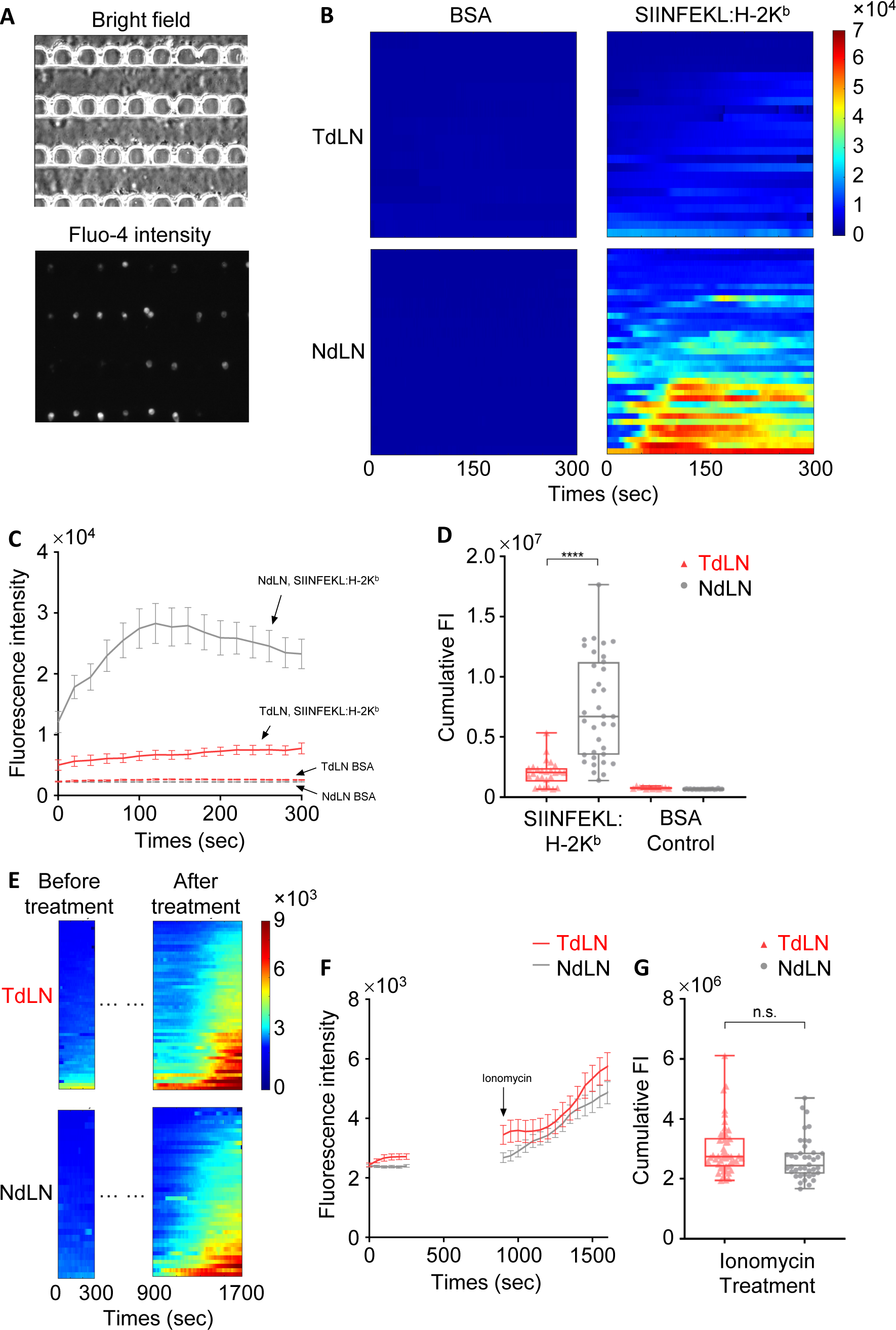
TdLN suppressed calcium response of CD8^+^ T cells to antigen stimulation. **(A)** Bright-field (*top*) and fluorescence (*bottom*) photomicrographs of the microfluidic cell trap array device used for the real-time calcium flux experiments. **(B)** Heatmaps of single-cell real-time Fluo-4 fluorescence intensity (FI) of the CD8^+^ T cells from TdLN (*top row*) or NdLN (*bottom row*) of an OT-I mouse in d7 post tumor implantation (representative results from three independent experiments). The inner surface of the cell traps was coated with either BSA (*left column*) or SIINFEKL:H2-K^b^ (*right column*). For each cell, time zero was the moment when the cell was trapped by a well, and each streak indicates the real-time Fluo-4 intensity for a tracked cell. Streaks were ranked in ascending order of cumulated FI over 300 sec. **(C)** Mean FI of the real-time calcium flux over all the cells from (*B*). **(D)** Cumulated FI over 300 s for each cell from (*B*). **(E-G)** Calcium response to perfusion of 1 μM ionomycin of CD8^+^ T cells from TdLN or NdLN of an OT-I mouse in d7 post tumor implantation (representative results from two independent experiments). Data presentation are parallel to (*B-D*), showing heatmaps (*E*), associated mean FI vs time (*F*) and cumulated calcium intensity (*G*) after ionomycin administration. Analyzed CD8^+^ T cells were recovered from TdLNs vs NdLNs of d7 B16F10 melanoma-bearing animals as depicted in Fig. 3A. Data are presented as mean ± SEM in (*C*) and (*F*), individual cells (points), mean (line) ± 75/25% (boxes) and max/min (whiskers) in (*D*) and (*G*). Statistical comparisons by Mann-Whitney test, ****p<0.0001, n.s.: p≥0.05.

Despite cell-to-cell variation, the overall calcium response to SIINFEKL:H2-K^b^ was much weaker for CD8^+^ T cells from TdLNs than NdLNs of the same tumor-bearing animal (Fig. 6B, right panels). This can also be seen in the averaged calcium fluorescent intensity (FI) vs time (Fig. 6C) and the FI accumulation over time for all individual cells (Fig. 6D), both of which were much lower for CD8^+^ T cells from TdLNs than NdLNs. As a positive control, CD8^+^ T cells from both tissue sources on BSA surfaces were stimulated by ionomycin, which showed robust and rapid calcium signals (Fig. 6E), displaying high levels of calcium signals without compartmental difference (Fig. 6F-G). Thus, rather than a TdLN-induced defect in the T cell’s intrinsic ability to flux calcium, the diminished calcium responses of CD8^+^ T cells from TdLNs is specific to defects in their capacity to recognize and respond to antigen. Together, these data indicate the reduced downstream signaling following TCR triggering in CD8^+^ T cells freshly isolated from the TdLN relative to NdLN.

### TdLN suppresses TCR 2D affinity and function of antigen-experienced CD8^+^ T cells

In the preceding sections, an “antigen-inexperienced” model where CD8^+^ T cells isolated from OT-I mice bearing B16F10 melanoma without the prior encountering of its cognate antigen *in vivo* was used to examine the TME effects *in vitro* for binding to (Figs. 3 and 4), pulling on (Fig. 5), and fluxing calcium on (Fig. 6) OVA pMHC. Whether the TME exerted similar effects on “antigen-experienced” CD8^+^ T cells was also explored using three models of *in situ* antigen recognition (Fig. 7A, D and G). In the first model, CD8^+^ T cells from CD45.2^+^ OT-I donor mice were adoptively transferred into CD45.1^+^ C57BL/6 recipient animals co-implanted in the lateral dorsal skin with B16F10 cells (Tumor) or left untreated (Naïve). Three days later, animals were immunized with Ag-NPs and Toll-like receptor ligand CpG-NP into the forelimb skin, a regimen we have previously shown to elicit robust CD8^+^ T cell immunity functional in mediating control of B16F10 melanomas (*37*). After 7 days post adoptive transfer (4 days post immunization), donor cells were harvested from the spleens and LNs and analyzed by FACS for activation markers and MAF for 2D affinity (Fig. 7A). The immunization step caused a population of CD8^+^ T cells to experience Ag stimulation *in vivo*. Whereas only small percentage of endogenous CD8^+^ T cells of the tumor-naïve CD45.1^+^ recipient mice isolated from the spleen and LN were CD44 positive, an indicator of activation by antigen (Fig. 7B, green), nearly all donor cells (CD45.2^+^) from both the spleen and LN of the recipient animals were CD44 positive (Fig. 7B, red). MAF experiments found lower OT-I TCR–SIINFEKL:H2-K^b^α3A2 2D affinity of donor cells from TdLNs than spleens of d7 B16F10 melanoma bearing animals; however, no such difference was found in tumor-naive recipient animals (Fig. 7C). This data indicates that TME locally suppresses TCR 2D affinity of CD8^+^ T cells primed by antigen *in vivo*.

**Fig. 7.**
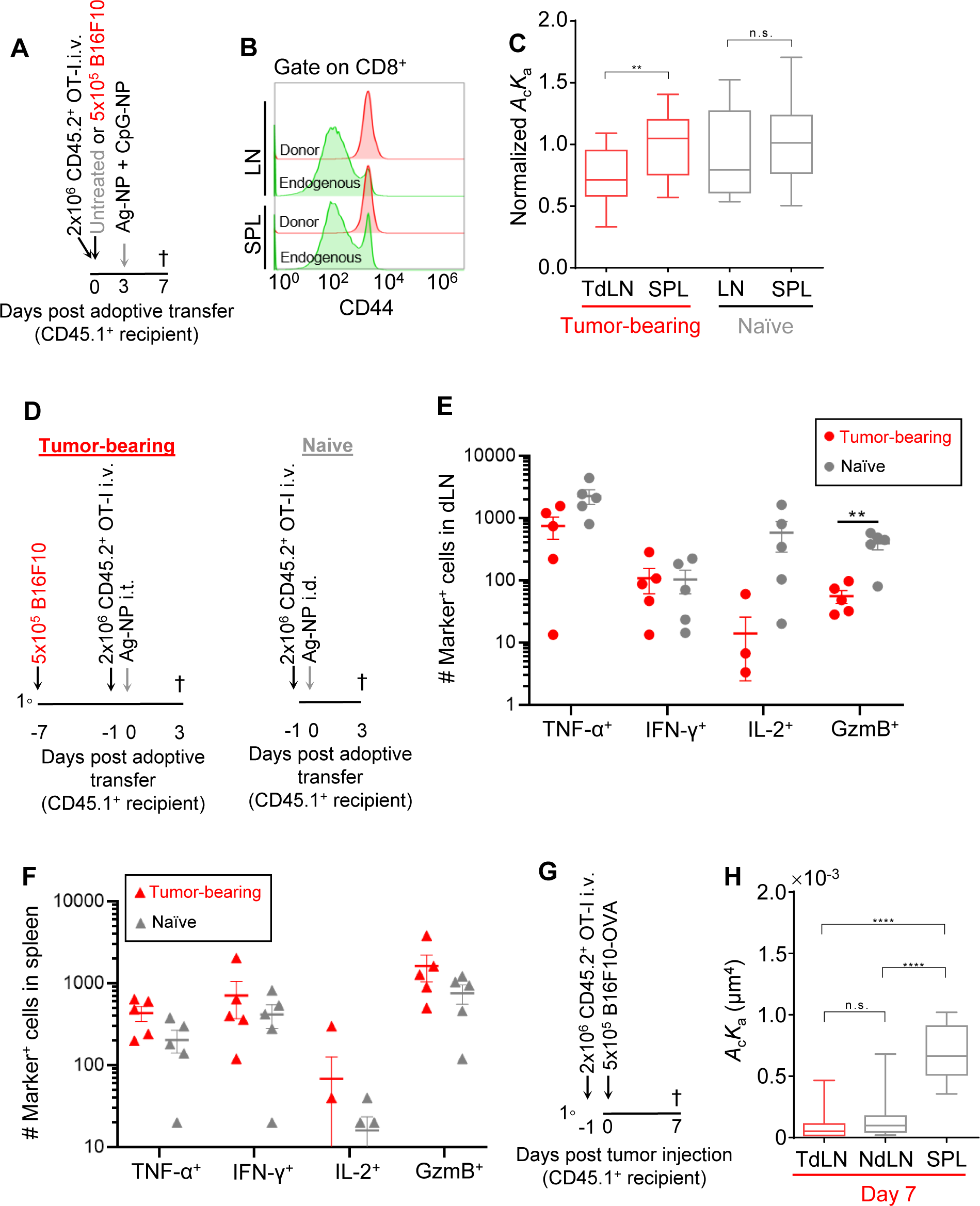
TdLN suppressed TCR–pMHC 2D affinity of antigen-experienced CD8^+^ T cells associated with altered profiles of activation and function. **(A)** Animal model used in (*B*-*C*). CD45.1^+^ C57BL/6J recipients implanted with B16F10 melanoma (Tumor) or left untreated (Naïve) were adoptively transferred with splenic CD8^+^ T cells from CD45.2^+^ OT-I mice at d0, immunized with Ag-NP and CpG-NP at d3, and sacrificed at d7 to isolate donor T cells from spleen and TdLNs (for Tumor-bearing mice) or LNs (for Naive mice) for analysis. **(B)** Comparisons of fluorescent histograms of CD44 expression of CD45.2^+^ (donor) and CD45.1^+^ (endogenous) CD8^+^ T cells from LNs and spleens in response to vaccination. Representative histogram of one sample from each group. **(C)** Comparisons of OT-I TCR–SIINFEKL:H2-K^b^α3A2 2D affinities of CD8^+^ T cells from spleen and TdLN of tumor-bearing recipients (red) and tumor-lacking recipients (gray), divided by the mean 2D affinity from the spleen of the same animal for each group to normalize variations between animals. Data presented as mean ± 75/25% (box) and max/min (whiskers) were pooled from two independent sets of experiments to acquire sufficient number (17–20) of T cells per condition to allow for reliable statistic comparison (Mann-Whitney test). **(D)** Animal model used in (*E*) and (*F*). *Left*, CD45.1^+^ C57BL/6J recipient animals were implanted with B16F10 melanoma at d-3. *Left and right*, donor splenic CD8^+^ T cells from CD45.2^+^ OT-I mice were adoptively transferred at d-1, and tumor-bearing (Tumor) or tumor-naïve (Naïve) animals were challenged with Ag-NP at d0 either i.t. or i.d., respectively, and animals were sacrificed at d3 to isolate donor T cells from TdLN and spleen (for tumor-bearing mice) or LN and spleen (for control mice) for analysis. **(E, F)** Comparisons of responses to *ex vivo* restimulated by SIINFEKL of donor cells harvested from TdLN vs LN (*E*) and spleens (*F*) of respective tumor-bearing vs -naïve animals by the number of cells expressing the indicated phenotypical markers. Each point represents data from an individual mouse. Data is the mean ± SEM. n=5. **(G)** Schematic of the model used in (*H*). Donor splenic CD8^+^ T cells from CD45.2^+^ OT-I mice were adoptively transferred d-1 into CD45.1^+^ C57BL/6J recipient animals implanted with B16F10-OVA B16F10-OVA tumors on d0. Donor T cells from spleen and TdLNs (for Tumor-bearing mice) or LNs (for Naive mice) were isolated from animals d7 for analysis **(H)** Comparisons of OT-I TCR–SIINFEKL:H-2K^b^α3A2 2D affinities of CD45.2^+^CD8^+^ donor T cells harvested from the indicated tissue compartments of the CD45.1^+^ C57BL/6J recipients bearing B16F10-OVA tumors. n=16-20 T cells pooled from repeated independent experiments. *indicates significance by two-way ANOVA with Tukey’s post-hoc test (*E*, *F*), or Mann-Whitney test (*C*, *H*).

The functional responses of CD8^+^ T cells that had previously experienced antigen to antigen re-exposure was assessed by the second model (Fig. 7D). Cell suspensions harvested from various tissues of tumor-bearing or -naïve CD45.1^+^ C57BL/6 recipient animals that had received adoptive transfer of CD8^+^ T cells from CD45.2^+^ OT-I mice and subsequently injected with SIINFEKL-NPs into the d7 B16F10 melanoma or naïve skin, respectively, were restimulated *ex vivo* using SIINFEKL peptide. In response to antigen restimulation, fewer donor cells from TdLNs of tumor-bearing mice than LNs of tumor-naïve mice produced TNF-α, IL-2, and GzmB, although similar numbers of cells produced INF-γ were found in the same two sites (Fig. 7E), implying decreased functional responsiveness by T cells in the tumor-draining vs naïve LN microenvironment. By comparison, donor cells recovered from spleens of tumor-bearing vs -naive animals exhibited no differences in TNF-α, IFN-ɣ, IL-2, and GzmB producing cells (Fig. 7F). This indicates that TME locally suppresses functional responses of antigen-experienced CD8^+^ T cells to antigen restimulation.

TCR 2D affinities were compared by the MAF experiments using the third model (Fig. 7G). CD8^+^ T cells from CD45.2^+^ OT-I mice were adoptively transferred into CD45.1^+^ C57BL/6 mice one day prior to implantation with B16F10-OVA melanoma cells. Donor cells isolated from TdLNs of recipients bearing d7 B16F10-OVA tumors exhibited a reduced TCR–pMHC 2D affinity compared to those from the spleen (Fig. 7H), consistent with previous results when the B16F10 melanoma implanted in the recipients was not conjugated with OVA antigen and hence the recovered donor cells were antigen-inexperienced. However, donor T cells harvested from NdLNs also displayed a lower TCR–pMHC 2D affinity than those harvested from spleens of the same recipient (Fig. 7H), in contrast to antigen-naïve models. Collectively, the results showed that the TME impairment of antigen recognition impacts CD8^+^ T cells not only at naïve stages but also after *in vivo* antigen activation.

## Discussion

Despite a functional systemic immunity, the melanoma TME is known to suppress its effects to allow disease progression (*55*). In the workhorse model of the melanoma immunotherapy field implemented in this work, antigen-induced activation and anti-tumor responses of CD8^+^ T cells *in vivo* was likewise suppressed by the TME. Whereas many cell intrinsic and extrinsic factors contribute the said suppression (*4, 5, 56*), our data reveal a new defect caused by the TME – the impaired T cell antigen recognition, manifested a reduced TCR–pMHC 2D affinity and bonds pulled by >4.7 pN T cell-generated forces measured *ex vivo.* This was further correlated functionally with a lower intracellular calcium signaling induced by *in vitro* antigen stimulation. These findings collectively indicate the impaired antigen recognition as a novel mechanism of T cell dysfunction in the TME. Interestingly, this newly identified effect of TME suppression of TCR antigen recognition may explain two puzzling observations of a recent study of adoptive cell therapy (ACT) of tumor patients (*57*). The authors reported that CD8^+^ T cells that were transfected with the same TCRs specific to tumor neoantigen, expanded in vitro, and used in ACT to treat cancer patients were more effective if 1) the original T cells were isolated from healthy individuals than from the patients themselves and 2) the original T cells were isolated from peripheral blood lymphocytes than from TILs of the same patient (*57*). The explanation based on the present work could be that the latter T cells had been subjected to the TME suppression whereas the former T cells had not in both cases.

The finding that CD8^+^ T cells from d3-d11 tumors and TdLNs (collectively referred to as TME) exhibit reduced *in situ* affinity of TCR for specific pMHC is particularly intriguing. Previous studies have shown correlation of 2D TCR–pMHC affinity with T cell activation and functions using altered peptide ligands (*17–23*). Unlike 3D affinity measured using soluble proteins, 2D TCR–pMHC affinity can be altered by perturbations of the T cell cellular environment (*17, 24, 25*). Of particular relevance to the present work, the P14 TCR–gp33:H2-D^b^ 2D affinity of adoptively transferred CD8^+^ T cells was found to be higher when donor cells were isolated from splenic red pulp than white pulp of C57BL/6 recipients post lymphocytic choriomeningitis virus infection. The increased antigen recognition of red pulp-derived CD8^+^ T cells correlates with a more efficient target cell killing *in vitro* and improved viremia control *in vivo*, whereas the decreased antigen recognition of white pulp-derived CD8^+^ T cells anti-correlates with their enhanced ability to differentiate to memory cells (*25*). Like the previous studies (*25*), differences in 2D affinity were not detectable by saturating pMHC tetramer staining, supporting the contention that 2D measurements made herein provide greater power to distinguish functionally relevant differences in binding propensities of TCR–pMHC interactions.

The TCR–pMHC–CD8 trimolecular avidity was also reduced by the TME, yet the CD8 affinities for both free MHC and TCR-bound MHC were not, isolating the TME-caused defects to the TCR itself. T cells can be activated by anti-TCR antibodies, yet the 2D binding of an anti-TCR antibody to TCR was not impacted, highlighting the specificity of the TME suppression on the ability of TCR to recognize antigen instead of being recognized as an antigen.

T cells spontaneously sample and exert forces on pMHC-coated surfaces. Such forces are transmitted through the TCR–pMHC complexes and thought to play a key role in antigen discrimination (*47, 48*). T cell spreading and pulling on pMHC was reduced by the TME, likely caused in part by the TME-suppressed TCR–pMHC and TCR–pMHC–CD8 interactions, because neither T cell spreading/pulling on anti-TCR antibody nor TCR–anti-TCR antibody interaction was suppressed by the TME. Reduced T cell spreading and pulling may be caused by the TME impairment of not only the recognition of, but also the response to, antigen by the T cell, which includes T cell signaling. This contention is supported by the results that only the calcium signaling of T cells stimulated by pMHC but not by ionomycin was reduced by the TME. Thus, the MTP experiments provide two additional readouts for the TME impairment of TCR recognition of, and activation by, antigen pMHC.

Chronic tumor antigen stimulation leading to T cell exhaustion is a well-known contributor to T cell dysfunction in the TME (*58*). However, we found that TME impaired antigen recognition by not only antigen-experienced but also antigen-inexperienced CD8^+^ T cells. In the absence of OVA-expression by tumor cells, antigen-naïve donor OT-I CD8^+^ T cells suppressed by B16F10 melanomas in recipient mice were not exhausted as evidenced by the low expression of PD-1. After isolation from the TME, these antigen-naïve cells encountered antigen the first time during the measurements for TCR–pMHC 2D affinity, endogenous force pulling on the TCR or calcium response, or during the *ex viv*o stimulation for functional assays. Since any donor T cells from any TCR transgenic mice can first be adoptively transferred to antigen-free tumor-bearing recipient mice and later be probed by the pMHC specific for this TCR to observe the TME effects, such suppressive effects must exert on all T cells of any antigen specificity. Thus, tumors can suppress the antigen recognition and function of T cells that do not recognize and react against the tumor itself, and this mechanism is independent of the T cell exhaustion due to chronic tumor antigen stimulation. This implies that the TME suppresses local CD8^+^ T cell immunity in general, not just that against tumor.

As a first step toward elucidating the mechanism underlying the TME suppression of T cell recognition of and response to antigen, we tested the role of TGF-β, a TME-associated immunosuppressive factor. The localized impairment of T cell antigen recognition was partially restored by inhibiting the TGF-β signaling. This indicates that, in addition to its other known effects in TME (*4, 5, 56*), TGF-β also negatively regulates all CD8^+^ T cells’ TCR–pMHC 2D affinities regardless of the TCR specificity. This agrees with our previous results that *in vivo* inhibition of TGF-β activity eliminates the differences in the TCR–pMHC 2D affinities between CD8^+^ T cells from splenic red and white pulps of virus infected CD8^+^ T cells and that *in vitro* treatment with recombinant TGF-β reduces TCR–pMHC 2D affinity (*25*). As the tumor progresses, TGF-β in the TME accumulates in the TdLN, which can affect any peripheral T cells trafficking through the TdLN. Comparing to other identified suppressive factors causing T cell tolerance, TGF-β is antigen-independent, unlike chronic tumor-antigen stimulation; and it does not require direct cell-cell contact between an immunosuppressive cell and the T cell, unlike nitration of tyrosine in TCR by MDSCs (*59*).

Together, our data uncover impaired T cell recognition of and response to antigen as a novel mechanism of T cell dysfunction in the TME. This mechanism is independent of T cell exhaustion induced by chronic tumor antigen stimulation and affects both antigen-inexperienced and antigen-experienced T cells irrespective of their antigen specificity. More importantly, the impaired T cell antigen recognition is a cause and an indicator of TME induced T cell dysfunction, which can be regulated by remodeling the soluble factor TGF-β in the TME. Our findings thus have implications for future studies seeking to identify TME extrinsic factors that suppress T cell function through altered TCR–pMHC 2D affinity and responses. Our results also implicate that the alteration of antigen recognition by CD8^+^ T cells can potentially serve as an additional metric for characterizing the TME and its amenability to immunotherapy.

## Methods

### Cell culture

B16F10 or B16F10-OVA mouse melanoma cells were maintained in culture in Dulbecco’s Modified Eagle Medium (DMEM, Gibco, Thermo Fisher Scientific, Inc., Waltham, MA) with 10% heat-inactivated fetal bovine serum (FBS, Gibco, Thermo Fisher Scientific, Inc.) and 1% penicillin/streptomycin/amphotericin B (Life Technologies, Carlsbad, CA, USA). Cells were passaged at ∼80% confluency and maintained at 37 °C with 5% CO_2_ in a standard cell incubator.

### Animal tumor models

CD45.2^+^ OT-I transgenic mice were obtained from Charles River Laboratory (Lyon, France) and bred in house at the Georgia Institute of Technology. C57BL/6 and CD45.1^+^ C57BL/6 mice were purchased at 6 weeks (wk) of age from the Jackson Laboratory (Bar Harbor, ME, USA). All protocols were approved by Georgia Tech’s Institutional Animal Care and Use Committee (IACUC) and have been previously described (*26, 38*). For tumor-bearing cohorts, 0.5 × 10^6^ melanoma cells were intradermally injected into the left dorsal skin of 6-8 wk old mice. Tumor dimensions were measured with calipers in three dimensions and reported as ellipsoid volume.

### Flow cytometry

Axillary and brachial LNs were pooled and incubated with 1 mg/mL Collagenase D (Sigma-Aldrich) in Dulbecco’s phosphate buffered saline (D-PBS) with calcium and magnesium for 75 min at 37 °C, passed through a 70 µm cell strainer (Greiner Bio-One, Monroe, NC, USA), washed, and resuspended in a 96-well U-bottom plate (VWR International, Inc.) for staining. Spleen capsules were disrupted using 18G needles (Beckton Dickinson, Franklin Lakes, NJ), and pushed through a 70 µm cell strainer, then pelleted and resuspended in RBC lysis buffer (Sigma-Aldrich) for 7 min at room temperature, diluted with D-PBS, washed, and 5% of the spleen plated in a 96-well U-bottom plate. Tumor samples were incubated with 1 mg/mL Collagenase D (Sigma-Aldrich) in D-PBS for 4 h at 37 °C, passed through a 70 µm cell strainer, washed, and 1-20% (based on tumor volume) plated in a 96-well U-bottom plate for staining. All antibodies for flow cytometry were obtained from Biolegend, Inc. (San Diego, CA, USA). Cells were blocked with anti-mouse CD16/CD32 (clone: 2.4G2) (Tonbo Biosciences, San Diego, CA, USA) for 5 min on ice and washed. Samples were then stained using fixable viability dye Zombie Aqua (1:100, Biolegend) for 30 min at room temperature before quenching with 0.1% bovine serum albumin (BSA) in D-PBS (flow cytometry buffer). Antibodies were prepared in flow cytometry buffer at the following dilutions based on preliminary titrations: PerCP anti-mouse CD3 (1:40), APC-Cy7 anti-mouse CD4 (1:640), FITC anti-mouse CD8 (1:320), PE-Cy7 anti-mouse CD25 (1:100), and APC anti-mouse PD-1 (1:80) to assess T cells in Fig. S1; APC-Cy7 anti-mouse CD45 (1:160), PerCP anti-mouse F4/80 (1:20), PE anti-mouse CD169 (1:50), APC anti-mouse Gr1 (1:80), PE-Cy7 anti-mouse CD11c (1:80), BV421 anti-mouse PD-L1 (1:40), FITC anti-mouse B220 (1:400), and AF700 anti-mouse CD11b (1:80) to assess pan immune cells in Fig. S1; PerCP anti-mouse CD69 (1:80), BV711 anti-mouse CD45.2 (1:80), BV605 anti-mouse CD3 (1:100), APC-Cy7 anti-mouse CD8 (1:40), and BV786 anti-mouse PD-1 (1:80) for intracellular cytokine staining in Figs. 2B-D and 7E-F; and PE anti-mouse CD45.2 (1:80), AF700 anti-mouse CD25 (1:100), BV785 anti-mouse PD1 (1:80), AF647 anti-mouse CXCR5 (1:200), PerCP anti-mouse CD3 (1:40), APC-Cy7 anti-mouse CD8 (1:40), PE-Cy7 anti-mouse CD39 (1:20), and BV421 anti-mouse CD44 (1:20) for T cell phenotyping in Fig. 1. Cells were fixed in 2% paraformaldehyde (VWR International, Inc.) to assess pan-immune cells in Fig. S1. For nuclear staining, cells were incubated with FoxP3/Transcription Factor Fixation/Permeabilization solution (eBioscience, ThermoFisher, Inc.) for 60 min on ice in dark. Cells were then incubated with PE anti-mouse FoxP3 (1:20) for T cell phenotyping in Fig. 1 and Fig. S1 in FoxP3/Transcription Factor Fixation/Permeabilization buffer for 75 min on ice in dark. For *ex vivo* cytokine staining, cells were suspended in IC Fixation Buffer (eBioscience, Thermo Fisher, Inc.) for 60 min at room temperature in dark. Cells were then incubated with APC anti-mouse GzmB (1:40), PE anti-mouse IFN-γ (1:80), AF700 anti-mouse IL-2 (1:200), and PE-Cy7 anti-mouse TNF-α (1:80) in IC Permeabilization Buffer (eBioscience, Thermo Fisher, Inc.) for 60 min at room temperature in dark. Cells were then resuspended in FACS buffer (1% BSA in D-PBS) and kept at 4°C in dark until analyzed with a customized BD LSR Fortessa flow cytometer (BD Biosciences). Compensation was performed using ArC and UltraComp compensation beads (Thermo Fisher Scientific, Inc.) and data was analyzed using FlowJo software v9 and v10 (FlowJo, LLC, Ashland, OR, USA).

### CD8^+^ T Cell Isolation

OT-I animals were sacrificed by CO_2_ asphyxiation. For adoptive transfer, spleens were dissected and placed into sterile PBS. Spleens were digested using 18G needles (Beckon Dickinson), and pushed through 70 μm cell strainers, washing liberally with PBS. RBCs were lysed using AcK Lysis Buffer (Lonzo Group AG, Basel, Swirzerland), and washed with PBS. After counting, cells were resuspended in FACS buffer (1% BSA in PBS) at a concentration of 10^8^ cells/mL and 1 μL negative CD8^+^ T cell biotin-antibody cocktail (Biolegend)/million cells was added and allowed to incubate for 15 min on ice. Streptavidin-magnetic beads were added directly to this solution at a concentration of 1 μL/million cells and allowed to incubate for 15 min on ice. PBS +/+ was then added for a final volume of 2.5mL and the mixture placed in magnetic separators. The flow-through was then collected, counted, and resuspended. CFSE was added to the cells for 6 minutes, followed by quenching with ice cold RPMI medium containing >10% heat-inactivated FBS (Life Technologies). Purity, viability, and CFSE loading (if applicable) were confirmed using a customized BD LSR Fortessa flow cytometer prior to transfer. Cells were maintained in sterile conditions prior to transfer.

For the CD8^+^ T cell isolation in MAF, MTP, and microfluidic calcium assays, organs of OT-I mice were mechanically digested into cell suspension, and CD8^+^ T cells were negative purified from cell suspension with untouched CD8^+^ T cell isolation kit (Stemcell Technologies). For the isolation of donor CD45.2^+^ OT-I T cells from C57BL/6 CD45.1^+^ recipient animals, the selection was achieved by antibody labeled sorting based on congenic marker.

### Adoptive Transfer

Isolated CD8^+^ T cells were suspended in sterile saline at a concentration of 2 × 10^6^ cells/200µL. After isoflurane anesthesia, the hair covering the jugular vein of animals was removed using depilatory cream, cleaned using warm water and ethanol wipes, and suspended cells injected intravenously via the jugular vein.

### TGF-β signaling inhibition

The animals were injected with TGF-β RI Kinase Inhibitor SB431542 (Sigma-Aldrich) at 10 mg/kg in 100 μl DMSO intraperitoneally for continuous three days.

### Micropipette adhesion frequency assay

The micropipette adhesion frequency (MAF) assay for measuring receptor–ligand 2D affinity has been described (*17, 39*). Briefly, human RBCs were biotinylated with controled concentration of biotin-X-NHS linker (Millipore/Sigma). The biotinylated RBCs were sequentially incubated with saturating amount of streptavidin (ThermoFisher) and the biotinylated proteins of interest. An RBC coated with ligand (L) and a T cell expressing receptor (R) were respectively aspirated by two apposing glass micropipettes (Fig. 2F). The two cells were brought into 50 contact-and-retract cycles by one of the micropipettes mounted to a piezoelectric translater controlled by a computer. The adhestion frequency (*P*_a_) was calculated by number of adhesion events appeared in the total of 50 contacts, where the adhesion events were detected by the deflection of the soft RBC membrane during micropiette retraction. The averaged number of bonds formed per contact <*n*> is calculated as:

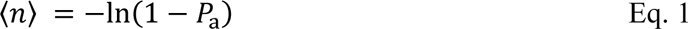

In the case of bimolecular interaction, the interaction kinetics is described by a probabilistic kinetics model:

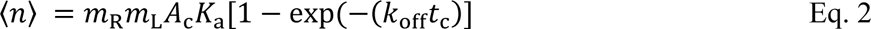

where *A*_c_ (in μm^2^) and *t*_c_ (in s) are contact area and time, respectively, controlled by the experimenter, *K*_a_ (in μm^2^) and *k*_off_ (in s^-1^) are 2D affinity and off-rate, respectively, *m*_R_ and *m*_L_ (in μm^-2^) are molecule densities of the receptor and ligand, respectively, determined from comparing the mean fluorescence intensity (MFI) of saturating amount of antibody staining of targeted proteins, to those of calibration beads (BD Biosciences). *P*_a_ reaches steady-state at *t*_c_ ≫ 1/*k*_off_ when exp(-*k*_off_*t*_c_) → 0, <*n*> → *m*_R_*m*_L_*A*_c_*K*_a_. Based on previous study (*17*), *P*_a_ reached plateau when *t*_c_ > 1 s for the OT-I TCR–SIINFEKL:H2-K^b^α3A2 interaction. Therefore, we report the values of effective 2D affinity *A*_c_*K*_a_ (in μm^4^) calculated from the equation below using *P*_a_ measured at 2 s.

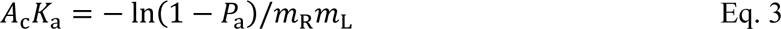

In the case where CD8^+^ OT-I T cells was allowed to interact with RBCs bearing SIINFEKL:H2-K^b^, <*n*> includes three species of bonds, TCR–pMHC and CD8–MHC bimolecular bonds and TCR–pMHC–CD8 trimolecular bonds (*18, 41, 48*). Neglecting the CD8–MHC bond species because of its orders of magnitude lower affinity than those of the other two bond species, we can express the average number of total bonds at steady-state by (*42*):

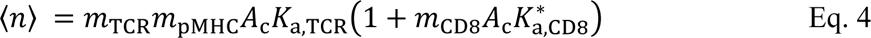

where 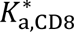 is the 2D affinity for CD8 to bind MHC pre-bound by TCR and *m*_CD8_ is the site density of CD8. 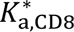 can solved from Eq. 4:

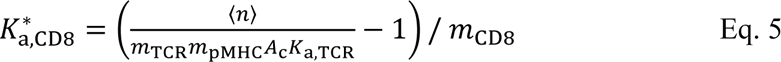

### DNA-based molecular-tension probe

The DNA-based molecular tension probes (MTP) were assembled as previously described (*47, 48, 60*). Briefly, three strands of oligonucleotides, the 4.7 pN hairpin strand, the Cy3B strand with biotin on the opposite end of the Cy3B dye, and the Black Hole Quencher-2 (BHQ2) strand, were mixed and annealed at a ratio of 1.1:1:1.1, with additional BHQ2 strand added after the annealing to further lower the background. The DNA-based MTP was then immobilized onto gold nanoparticles irreversibly anchored on APTES-PEG modified glass surface (*47*). The surface was sequentially incubated with streptavidin and biotinylated pMHC ligands or antibodies to tag them to the MTPs through biotin-streptavidin coupling. Working solution, incubation time and washing buffer for each step were described previously (*47, 48, 60*). The MTPs was assembled on glass slides and ready for use.

The CD8^+^ OT-I cells were injected into the image chamber, allowed to settle and spread on the glass surface for 20 min, and imaged for 30 min with a PerkinElmer confocal microscope with TIRF mode (Hamamatsu sCMOS detector, Nikon Ti-E Microscope, 60× 1.49 TIRF oil objective). The Reflection interference contrast microscopy (RICM) image of a cell appeared as a dark area which was defined as the cell-surface contact area. The tension signal of a cell was measured as the mean Cy3B fluorescence intensity (FI) observed in the total internal reflection fluorescence (TIRF) mode. It was calculated by subtracting the surrounding background level FI from the original contact area FI, and averaging over the spreading area, except for the negative control in which the signal was normalized by the cell size detected on the bright field image, since no detectable cell-surface contact was formed.

### Microfluidic cell trap real-time calcium flux analysis

The microfluidic cell trap arrays have previously been described (*54*). Briefly, a layer of polydimethylsiloxane (PDMS) (10:1) was poured on the previously designed and fabricated 8-μm master mold, vacuumed overnight for debubbling, and baked at 70 °C for 2 h before being peeled off from the mold, cut out and punched the inlet and outlet holes for individual devices. For each PDMS block, the side with micro-channels was treated with oxygen plasma to let it firmly bond to a coverslip. The device was completed by connecting tygon tubing to the inlet and outlet on the PDMS. To functionalize the inner surface to be contacted by cells, the devices were washed with PBS, coated in turn with biotin-BSA for 1 h, streptavidin for 30 min, and biotinylated ligand (0.2 mg/ml) for 30 min at room temperature. To avoid unwanted transient stimulation of cells bumping into the coated surface and inlet filter pillars before entering the trapping zone, the inlet channels and filter was blocked with BSA during the biotin-BSA coating step, so the ligand would not be coated to these areas. The device was washed between each coating step by perfusing PBS.

CD8^+^ T cells were loaded with 5 μM of Fluo-4, AM calcium indicator (Thermo Fisher F14201) at 37 °C for 30 min. Calcium indicator loaded cells at 2 × 10^6^/ml in the imaging buffer (Hanks’ Balanced Salt Solution, or HBSS, with calcium/magnesium and 2% FBS, passed through a 0.45 μm filter) were stabilized for another 30 min at 37 °C. Cell suspension was aspirated into 10 μl pipettor tip and added into the microfluidic device through the inlet. To test the calcium response to soluble factors, calcium indicator-loaded CD8^+^ T cells were first trapped in the device wells, followed by perfusion of the soluble stimulator. The cell trapping and calcium signaling process was recorded using a PerkinElmer confocal microscope system (Hamamatsu EM-CCD detector, Nikon Ti-E Microscope, 20× 0.75 Air Objective, LiveCell temperature control incubator). The fluorescent signal was recorded continuously for 10 min at 2 frame per sec (fps) for the first 5 min and 1 fps for the rest time. Snap shots of bright field images were taken before and after the 10 min recording for the device and cell trap quality control. The experiments were performed at 37 °C.

Single-cell real-time calcium intensity was analyzed using fluorescent time series data stack. The analysis was similar to those described previously in principle, with the current version of program written in Fiji in house (available upon request). Briefly, the moment that a cell falls into the cell trap is defined as time zero for this cell. The program generates a mask over this cell and the MFI of this mask is tracked in the following frames. This analysis was applied to all cells appeared in the field of view, and then the background FI was subtracted from the MFI. The calcium signal level vs. time sequences for different cells were then realigned at time zero for display and sorting.

### Synthetic antigen system

Pyridyl-disulfide functionalized nanoparticles were prepared as previously described (*30, 32, 61*). Cysteine-modified SIINFEKL (CSIINFEKL) was dissolved in MilliQ water at 1 mg/mL and added 1:1 to 40 mg/mL PDS-NP. The disulfide displacement reaction proceeded overnight at room temperature with stirring after which SIINFEKL conjugated NP were separated from unreacted peptide by size exclusion chromatography using a CL-6B column. Fractions containing peptide were identified by reacting with fluorescamine, and PEG-containing fractions (NPs) were determined using an iodine assay. Fractions containing CSIINFEKL-NP (Ag-NP) were combined and concentrated to the appropriate dose using 30 kDa MWCO spin filters and sterilized by filtration through a 0.22 µm syringe filter. In select experiments, NPs were reacted with maleimide-AlexaFluor647 or maleimide-AlexaFluor700 during synthesis to render the NP fluorescent (*29–32*). Ag-, AF647-, or AF700-NP in sterile saline were injected intradermally in the center of the tumor or into the dermal layer of the skin (non-tumor-bearing animals) of C57Bl/6 mice.

### *Ex vivo* Restimulation

After cell isolation (as above, in flow cytometry), 30% of LN samples, 5% of spleen samples, or 5% of tumor cells were plated in a sterile 96-well U-bottom plate. 1 µg/mL SIINFEKL peptide in 200 µL IMDM media with 10% heat-inactivated FBS and 0.05 mM betamercaptoethanol (Sigma-Aldrich) were added to each sample and incubated for a total of 6 h at 37 °C with 5% CO_2_. Three hours into the incubation period, 50 µg/mL Brefeldin A (Sigma-Aldrich) was added to each sample. Cells were washed and stained for flow cytometry as above.

### Imaging by In Vivo Imaging System (IVIS)

Animals were injected intradermally in the left dorsal skin with AF647 conjugated NP as described above. 24 h after NP injection, animals were sacrificed via CO_2_ asphyxiation in accordance with AVMA and local IACUC guidelines. Animals were dissected to expose axillary and brachial LNs and imaged using a Perkins Elmer IVIS Spectrum CT (Waltham, MA). LNs were then dissected and placed on black plastic and imaged using a Perkin Elmer IVIS Spectrum CT.

### Tumor Resection

Animals were anesthetized using isoflurane in oxygen, and then given 1 mg/kg sustained release Buprenorphine and 5 mg/kg ketoprofen via intraperitoneal injection as analgesics. The animals were placed on a warming bed and a sterile, fenestrated drape, placed to expose only the tumor and surrounding skin. Povidone-iodine was applied to the skin three times to sterilize the surgical area. Sterile scissors were used to excise and remove the tumor, and sterile wound clips used to close the wound. The animal was monitored throughout recovery and returned to the cage. Animals were monitored every other day to ensure well-being and examine for infection or irritation surrounding the surgical site. Wound clips were removed 10 days post-surgery. All procedures were in accordance with AVMA and local IACUC guidelines.

### Statistical analysis

Data are presented as the mean accompanied by SEM or SD and statistics were calculated using Graphpad Prism 6, 7, and 8 software (Graphpad Software, Inc., La Jolla, CA, USA). Statistical significance was defined as p<0.05, 0.01, 0.05, and 0.001 respectively, unless otherwise specified.

## Acknowledgement

We thank B. Liu, Z. Li, and Y. Chen for contributing to the feasibility studies at the initial stage of this project. This work was supported by NIH grants U01CA214354, R01CA207619, T32GM008433, and T32EB021962. M.P.M. was supported by a National Science Foundation Graduate Research Fellowship.

## Author Contributions

C.Z. and S.N.T. conceived and supervised the project; Z.Y., M.J.O., J.L., F.Z., P.J., K.B, C.G., M.N.R., L.C., S. R.-C., N.A.R., M.P.M, and D.M.F. performed experiments and analyzed data; K.L. developed image analysis program; H.L. provided specialized devices; C.Z., S.N.T., Z.Y., and M.J.O. wrote the paper.

---

**Supplemental Fig. 1.**
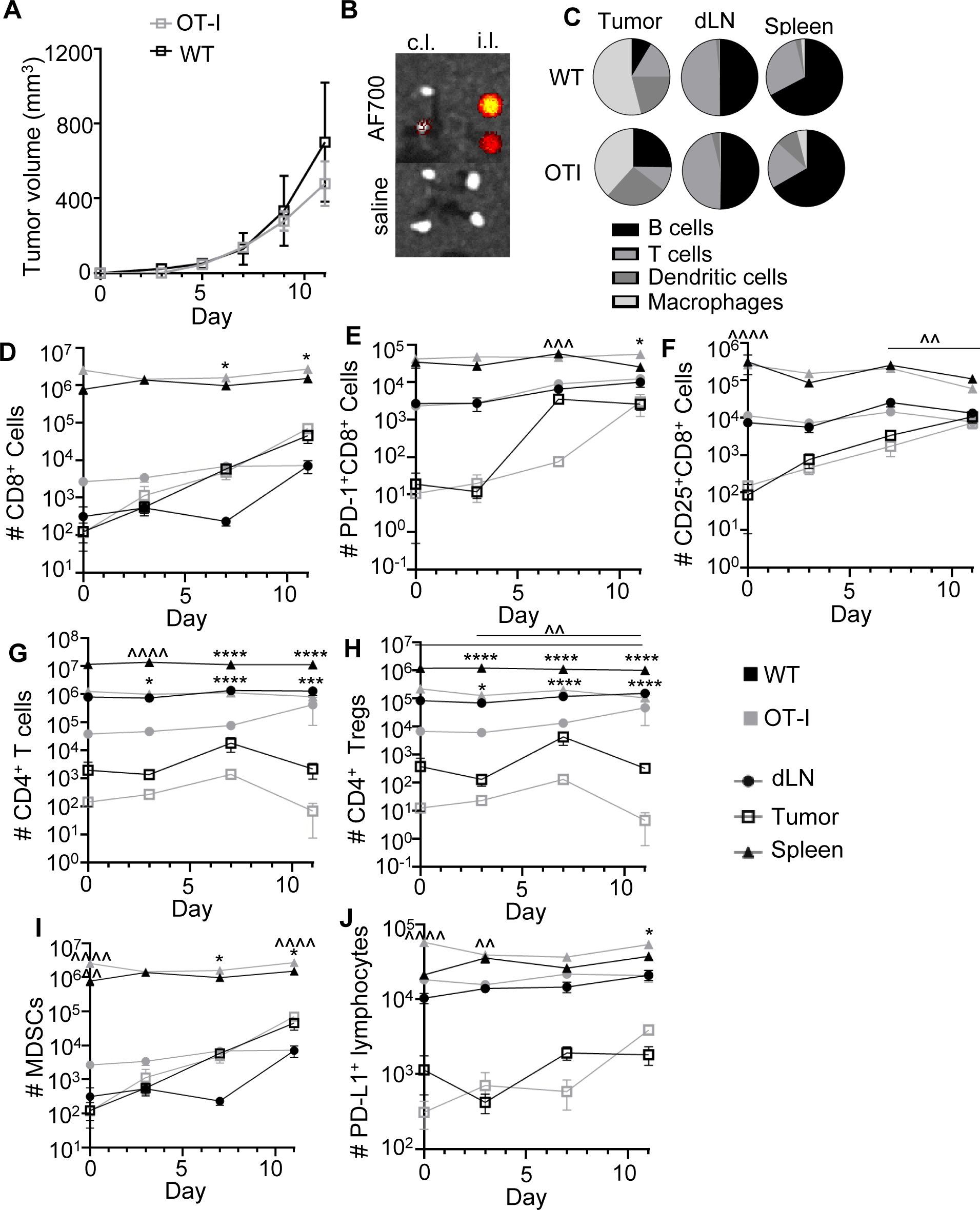
Tumor growth and immune phenotyping of B16F10 tumor growth in C57Bl/6 versus OT-I mice. **(A)** Volumes of B16F10 melanomas implanted in C57BL/6J and OT-I animals. **(B)** Identification of dLN by intratumoral injection of AF700-labelled lymph-draining particles (*top, right*) comparing to saline injection (*bottom*). **(C)** Major immune cell subtypes in animals bearing d7 B16F10. **(D-J)** Number of CD8^+^ T cells (*D*), PD-1^+^CD8^+^ T cells (*E*), CD25^+^CD8^+^ T cells (*F*), CD4^+^ T cells (*G*), Tregs (*H*), MDSCs (*I*), and PD-L1^+^CD45^+^ cells (*J*), in naïve, d3, d7, and d11 B16F10 tumors, dLNs, and spleens. “*” indicates difference between animal type by two-way ANOVA; “^” indicates difference between timepoints in each animal type (against all other timepoints if not specified). Data represent ± SEM (*A*, *D*-*J*). n=5.

**Supplemental Fig. 2.**
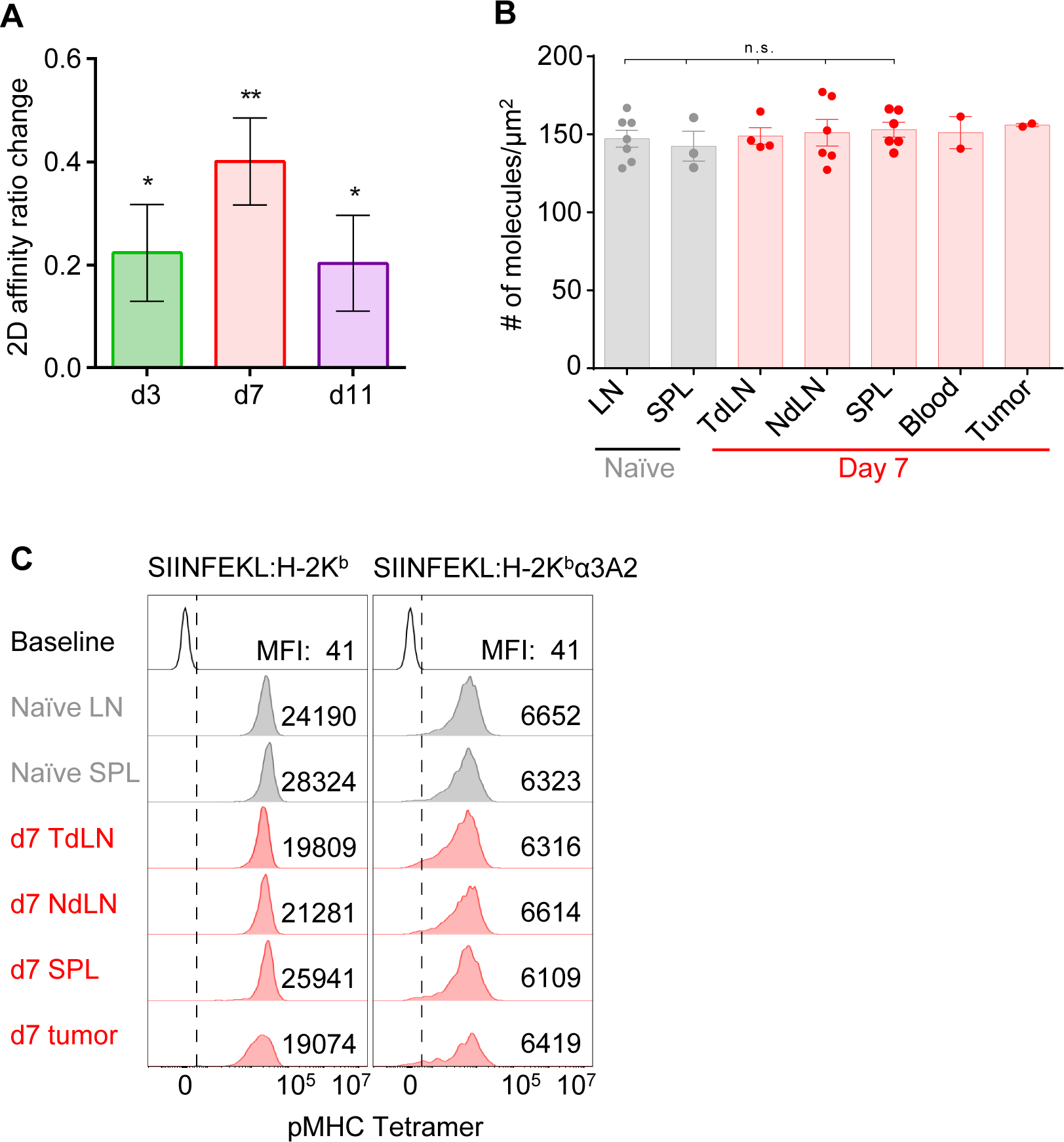
Assessment of CD8^+^ T cells from various tissue compartments of B16F10-bearing (red) and -lacking (gray) OT-I animals. **(A)** Ratios of OT-I TCR– SIINFEKL:H2-K^b^α3A2 2D affinity of CD8^+^ T cells from TdLN over that from NdLN in d3, d7 and d11 post tumor implantation. Data are presented as mean ± SD and assessed statistically by one sample t tests with hypothesis form zero. **(B, C)** TCR expression on CD8^+^ T cells from indicated tissue sources in mice bearing (red) or lacking (gray) a d7 tumor, shown as MFI of saturating concentration staining by an anti-TCR antibody (*B*) and FI histograms (curves) and MFI (numbers) of saturating concentration staining by SIINFEKL:H-2K^b^ (*left*) and SIINFEKL:H-2K^b^α3A2 tetramers (*right*) (*C*).

